# Mediator of DNA damage checkpoint 1 (MDC1) is a novel estrogen receptor co-regulator in invasive lobular carcinoma of the breast

**DOI:** 10.1101/2020.12.16.423142

**Authors:** Evelyn K. Bordeaux, Joseph L. Sottnik, Sanjana Mehrotra, Sarah E. Ferrara, Andrew E. Goodspeed, James C. Costello, Matthew J. Sikora

**Affiliations:** Dept. of Pathology, University of Colorado Anschutz Medical Campus; Biostatistics and Bioinformatics Shared Resource, University of Colorado Comprehensive Cancer Center; Dept. of Pharmacology, University of Colorado Anschutz Medical Campus

**Keywords:** Invasive lobular carcinoma, breast cancer, estrogen receptor, co-regulator, MDC1

## Abstract

Invasive lobular carcinoma (ILC) is the most common histological subtype of breast cancer, and nearly all ILC tumors express estrogen receptor alpha (ER). However, clinical and laboratory data suggest ILC are strongly estrogen-driven but not equally sensitive to anti-estrogen therapies. We hypothesized that ILC-specific ER transcriptional co-regulators mediate ER functions in ILC and anti-estrogen resistance, and profiled ER-associated proteins by mass spectrometry. Three ER+ ILC cell lines, MDA MB 134VI, SUM44PE, and BCK4, were compared to published data from ER+ invasive ductal carcinoma (IDC) cell lines, and we examined whether siRNA knockdown of identified proteins suppressed ER-driven proliferation in ILC cells. This approach found mediator of DNA damage checkpoint 1 (MDC1), a key tumor suppressor in DNA damage response (DDR), as a putative novel ER co-regulator in ILC. We confirmed ER:MDC1 interaction was specific to ILC cell lines versus IDC cells, and found MDC1 knockdown suppressed ILC cell proliferation and suppressed tamoxifen resistance in MDA MB 134VI. Using RNA-sequencing, we found that in ILC cells, MDC1 knockdown broadly dysregulates the estrogen-driven ER transcriptome, with ER:MDC1 target genes enriched for hormone-response-elements in their promoter regions. Importantly, our data are inconsistent with MDC1 regulating ER via MDC1 DDR and tumor suppressor functions, but instead suggest a novel oncogenic role for MDC1 in mediating ER transcriptional activity as a co-regulator. Supporting this, in breast tumor tissue microarrays MDC1 protein was frequently low or absent in IDC or ER-ILC, but MDC1 loss is rare in ER+ ILC. ER:MDC1 interaction and MDC1 co-regulator functions may underlie cell type-specific ER functions in ILC, and serve as important biomarkers and therapeutic targets to overcome anti-estrogen resistance in ILC.

## INTRODUCTION

Invasive lobular carcinoma (ILC) is the most common histological subtype of breast cancer relative to invasive ductal carcinoma (IDC, no special type). ILC accounts for 10-15% of new breast cancer diagnoses (∼40,000 new cases annually) in the United States, making ILC among the top 10 most frequently diagnosed cancers affecting women [1, 2]. Despite this prevalence and the unique clinical presentation of ILC [3, 4], only recently have clinical and laboratory studies focused on ILC begun to characterize ILC as a unique disease among breast cancers.

As ∼95% of ILC express estrogen receptor alpha (i.e., are ER+) and >80% classify as the hormone-dependent Luminal A molecular subtype [2], ILC is presumed to respond to anti-estrogens (tamoxifen or aromatase inhibitors, AIs); however, prospective studies of ILC outcomes have yet to be reported. In retrospective studies compared to IDC, patients with ILC have worse long-term outcomes, regardless of ER status. Reduced disease-free and overall survival for patients with ILC >5-10y post-diagnosis have been confirmed in studies in Europe (total n=18397, n=2033 ILC; [5–7]) and the US (NCI SEER; n≈800k; n≈85k ILC [8]). In SEER, ILC patients had decreased disease-free survival (IDC:ILC HR=0.809, p<0.0001) and overall survival (HR=0.775, p<0.0001) beyond 5 years post-diagnosis [8]. These data parallel the decades of recurrence risk faced by patients with ER+ cancer [9], and indicate that despite being largely ER+/Luminal A, current therapies are not curative for many patients with ILC.

Though nearly all patients with ILC are treated with anti-estrogens (since ∼95% are ER+), the efficacy of anti-estrogens for ILC is unclear. To address this, Metzger et al analyzed the BIG 1-98 trial (n=2923, n=324 ILC), which compared adjuvant tamoxifen to the AI letrozole [10]. Letrozole-treated patients with ILC had similar outcomes to IDC; however, tamoxifen-treated ILC patients had increased recurrence and poorer survival (8y recurrence 34%, vs 25% in IDC; 8y overall survival 74%, vs 84% in IDC). Poor outcomes and tamoxifen resistance were also reported for the ABCSG-8 trial [11]. Paralleling clinical data, we and others find ER+ ILC models are tamoxifen-resistant, including *de novo* ER partial agonism by anti-estrogens, including both tamoxifen and oral selective ER degraders [12–16]. Further, in cell line and *in vivo* ILC models, we found ER regulates hundreds of unique target genes via distinct ER DNA binding [12], and that ER regulates novel signaling pathways critical for endocrine response and resistance in ILC, such as cell-intrinsic Wnt signaling via WNT4 [12, 17–19]. Taken together, these data suggest ER has distinct function in ILC cells versus other breast cancer cells.

Transcription factors including ER control gene expression by binding DNA to recruit transcriptional machinery including co-regulator proteins [20]. Importantly, cell type-specific co-regulator functions can mediate cell type-specific transcription factor activity and responsiveness to selective ER modulators (SERMs) [21, 22], suggesting that co-regulator activity may mediate ILC-specific ER functions and SERM resistance. Consistent with this, data from the Cancer Genome Atlas (TCGA) also suggest that ILC is a unique context for ER function. In breast cancer cells, ER requires the pioneer factors FOXA1 and GATA3 to access DNA [23, 24], but mutations in these factors are differentially enriched in ILC versus IDC (among Luminal A tumors). *FOXA1* mutations are enriched in ILC vs IDC (7% vs 2%), while *GATA3* mutations are depleted in ILC vs IDC (5% vs 13%) [2]. The role of these mutations is unclear, but these data support that ILC-specific functions of ER-interacting proteins create a unique context for ER function in ILC cells.

Based on these observations, we hypothesized that unique ER co-regulators mediate distinct ER functions in ILC, and profiled ER interacting proteins by the immunoprecipitation/mass spectrometry (IP/MS) method RIME (<u>R</u>apid <u>I</u>P and <u>M</u>S of <u>E</u>ndogenous proteins [25]). RIME uses crosslinking with formaldehyde prior to IP to preserve protein complexes, and was developed to identify ER co-regulators [26]. This approach led us to identify MDC1 (mediator of DNA damage checkpoint 1) as a putative novel co-regulator of ER in ILC cells, where we found MDC1 is required for ER-driven proliferation and gene regulation. These data led us to further investigate the novel ER co-regulator functions of MDC1 in ILC cells.

## RESULTS

### RIME identifies ILC-specific ER-associated proteins and putative novel co-regulators

To identify ILC-specific ER co-regulator proteins, we profiled ER-associated proteins in ILC models using RIME [25, 26]). We performed ER RIME with ER+ ILC cell lines MDA MB 134VI (MM134), SUM44PE (44PE), and BCK4 in full serum treated with either vehicle (0.1% EtOH) or 1µM 4-hydroxytamoxifen (4OHT). MS analyses identified 416, 231, and 333 ER-interacting proteins in MM134, 44PE, and BCK4 respectively (**Figure 1A; Supplemental File 1**). Notably, few proteins specific to either vehicle or 4OHT conditions were identified (**Supplemental Figure 1**). To identify putative ILC-specific ER-associated proteins, we compared proteins identified in at least 2 ILC models (n=188) to ER RIME data from IDC cell lines MCF7 and ZR75-1 (union, n=713 [26]). This identified ILC-specific ER-associated proteins (n=115), and ER-associated proteins common to ILC and IDC cells (n=73) (**Figure 1B; Supplemental File 1**). We queried these proteins in the STRING database [29] (**Figure 1C-E**), and among ILC-specific ER-associated proteins this identified a network of epigenomic regulators and known co-regulators (e.g. DNMT1, KMT2D, BRD4, SAFB), and a DNA repair network (e.g. FEN1, POLD2) (**Figure 1C, E**). Among ER-associated proteins common to ILC and IDC cells, this identified a gene regulatory network of established ER co-regulators (e.g. EP300, GATA3, FOXA1, GREB1, NCOA3) (**Figure 1D, E**). These data show ER associates with a core network of common co-regulator proteins in ILC and IDC cells, but that ER also interacts with network of unique putative co-regulators in ILC cells.

**Figure 1.**
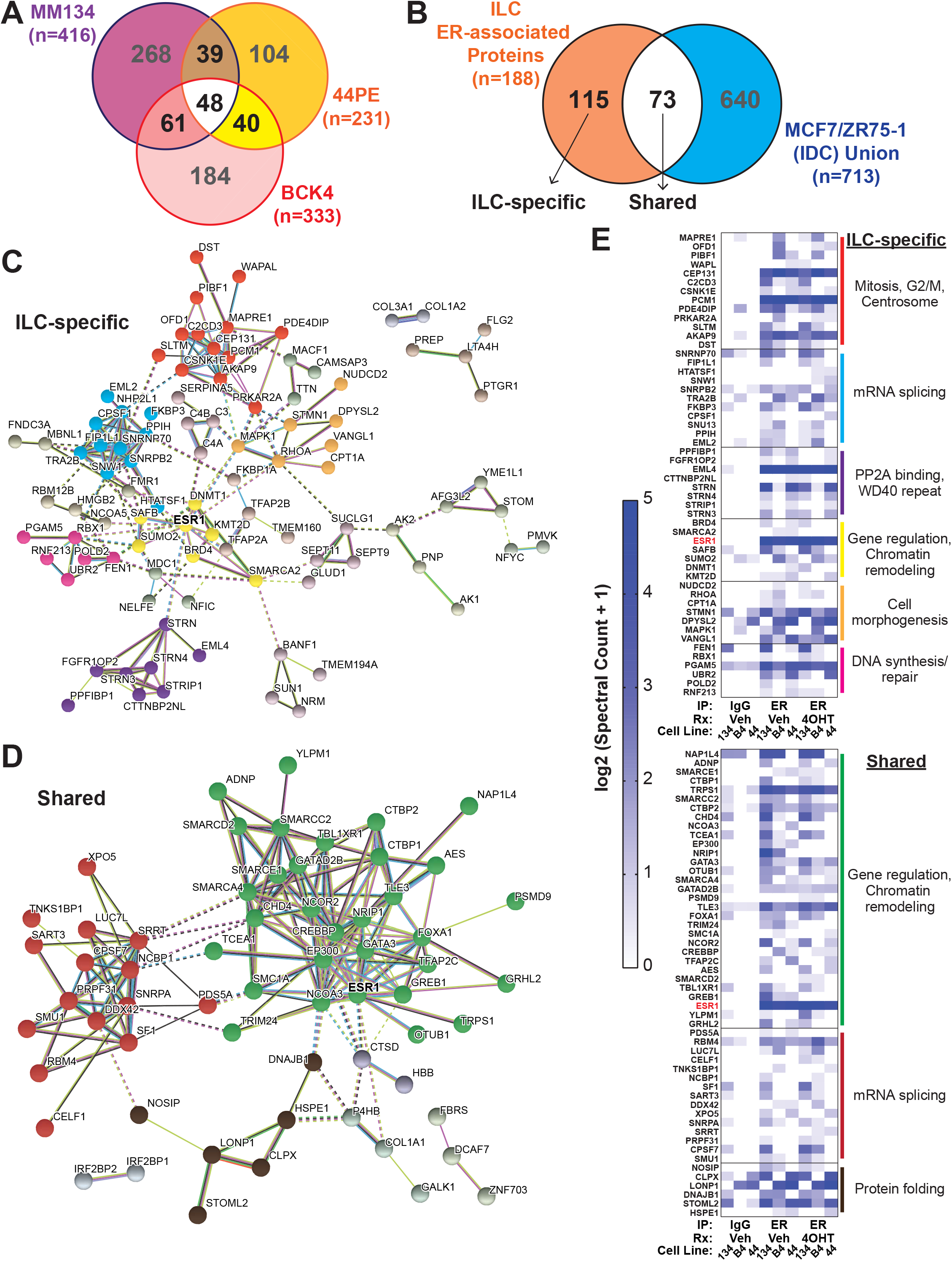
RIME identifies unique ER-associated proteins in ILC cell lines. **(A)**, ER co-IP/MS was performed using RIME as described in Materials and Methods using three ER-positive ILC cell lines. ER-associated proteins identified in at least 2 cell lines were used in subsequent analyses. **(B)**, Proteins identified in (A) were compared to ER RIME in IDC models MCF7 and ZR751 from Mohammed, 2013. **(C)**, ILC-specific ER-associated proteins (n=115; plus ER/ESR1, total n=116) were used for network analyses using the STRING database (v11.0; October 2019). Colored clusters were generated in STRING using MCL clustering. **(D)**, STRING network analysis as in (C) for ER-associated proteins common to ILC and IDC (n=73, including ER). **(E)**, Clusters from (C-D) with >5 members are highlighted; colored bars match circle colors in (C-D) network maps. Heatmaps show the mean spectral counts of technical duplicates for each protein, per cell line and treatment/IP condition. 134 = MDA MB 134VI; B4 = BCK4; 44 = SUM44PE. Functional notations for clusters listed at right are derived from gene ontology analyses using MSigDB/DAVID.

### Mediator of DNA damage checkpoint 1 (MDC1) is critical for ER function in ILC cells

We functionally profiled identified ER-associated proteins using an siRNA screen to determine how target knockdown affected ER activity in MM134 (ILC) cells, and hypothesized that knocking down ER co-regulators would suppress ER-mediated cell proliferation driven by E2 or tamoxifen [12]. We selected 133 proteins from RIME for screening, supplemented with co-regulators over-expressed in ER+ ILC versus ER+ IDC (n=25; n=10 other differentially expressed co-regulators were added *ad hoc)*. We also included transcription factors with binding motifs flanking an ER binding site for the ILC-specific ER target gene *WNT4* as factors with putative functions to regulate ER activity in ILC [17](n=31) (total n=199, **Supplemental File 2**). MM134 cells were hormone-deprived prior to siRNA transfection; 24hrs post-transfection, cells were treated with vehicle (0.01% EtOH), 100pM estradiol (E2), or 100nM 4OHT. Additionally, because androgen receptor (AR) can drive growth independently of ER in MM134 (**Supplemental Figure 2A**), we treated cells with synthetic androgen Cl-4AS-1 (100nM) for counter-screening to identify ER-specific co-regulators. Six days post-treatment, cell proliferation was assessed by dsDNA quantification, and we identified siRNAs that suppressed E2- and/or 4OHT-driven growth, but did not reduce growth in vehicle- and/or androgen-treated conditions (complete dataset in **Supplemental File 3**). Of note, siRNA targeting the ILC-specific ER-centered ‘gene regulation’ network (yellow in **Figure 1C/E**, e.g. KMT2D, BRD4) did not pass the latter criteria and suppressed cell number in vehicle conditions, suggesting a role for these proteins in ILC cell viability (**Supplemental Figure 2B**). Using these filters, we identified 82 targets for which knockdown suppressed ER-driven proliferation (i.e. via E2 or 4OHT) (**Figure 2A, Supplemental File 3**). siRNAs targeting ER itself (siESR1) and pioneer factor FOXA1 (siFOXA1) were among the top hits, acting as positive controls for the siRNA screen; few siRNAs showed differential suppression of E2-versus 4OHT-induced growth (**Supplemental Figure 2C**). Interestingly, siRNA targeting *MDC1* (siMDC1) was equivalent to siESR1 in suppressing both E2- and 4OHT-induced growth, and not affecting AR-driven growth, suggesting MDC1 is associated with ER function (**Figure 2B**). Based on this, we further examined MDC1 as a novel ER co-regulator in ILC cells.

**Figure 2.**
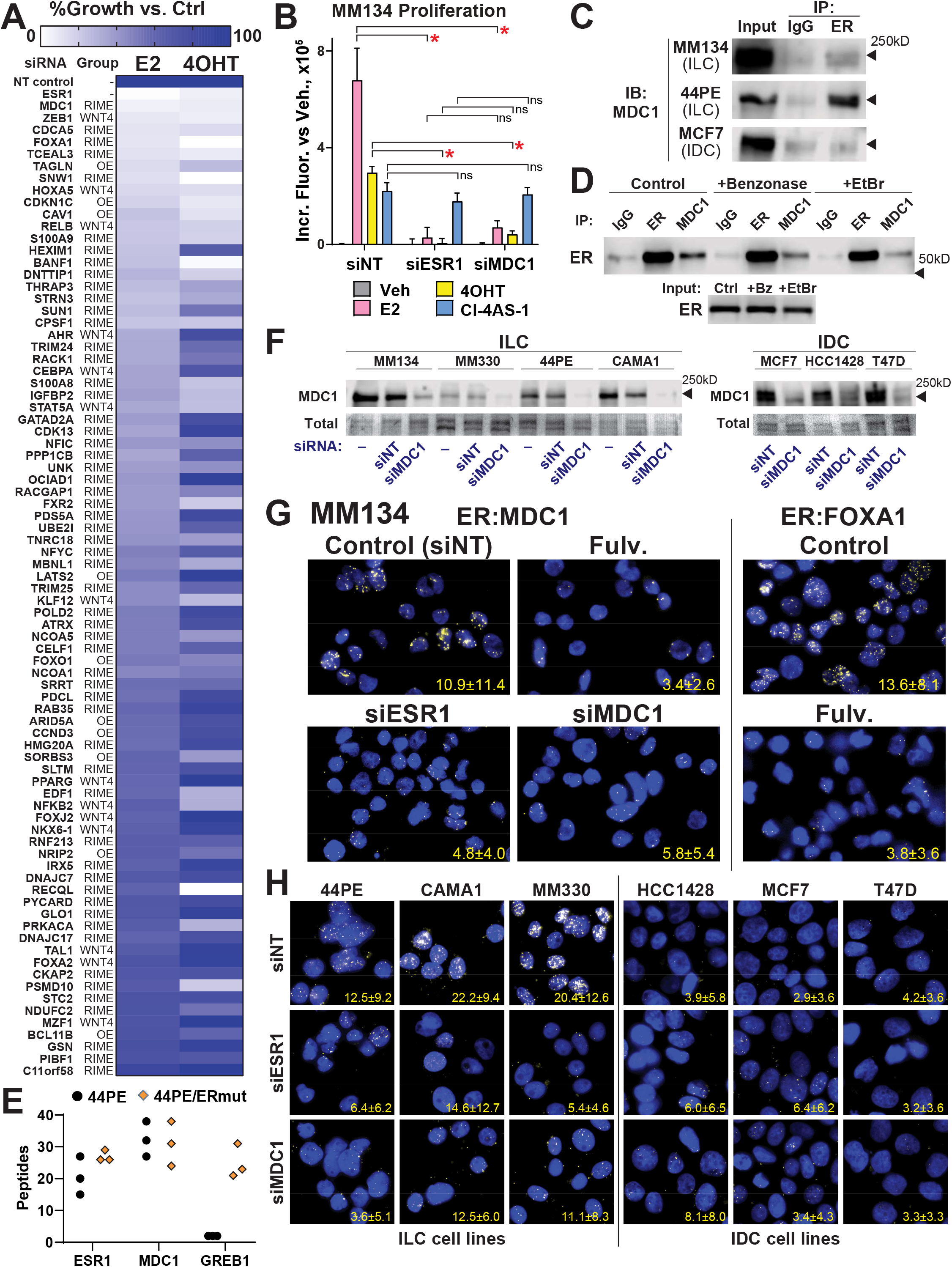
MDC1 is a novel ER-associated protein in ILC cells and is required for ER-driven proliferation. **(A-B)**, MM134 cells were hormone-deprived prior to siRNA transfection, then treated with vehicle (0.01% EtOH), 100pM E2, 100nM 4OHT, or 100nM Cl-4AS-1 (synthetic androgen). After 6d, proliferation was measured by dsDNA quantification. **(A)**, Growth vs mock siRNA control for E2 and 4OHT treatments. ‘Group’ represents context for gene inclusion in siRNA panel: RIME= identified by RIME; WNT4= transcription factor motif at ER binding site in *WNT4* gene; OE= over-expressed in ER+ ILC vs ER+ IDC. **(B)**, Data from screen in (A). *, p<0.05; n.s. = not significant; ANOVA w/ Dunnett’s multiple correction. **(C)**, Nuclei (input) from cells in full serum were extracted prior to co-IP; ER co-IP enriched MDC1 vs IgG in ILC cell lines but not MCF7 (IDC). **(D)**, Nuclear extracts were treated with benzonase (+Bz) or ethidium bromide (+EtBr) prior to co-IP to disrupt DNA structure and prevent DNA-dependent co-IP. MDC1 co-IP enriched ER vs IgG, reciprocal co-IP to (C). **(E)**, Peptide counts from Martin et al 2017, points represent biological triplicate RIME samples. **(F)**, Breast cancer cells were transfected with siNT or siMDC1 48hrs prior to lysate harvest for immunoblot. -, mock transfection (reagent only). Single v double bands at ∼250kD are related to polyacrylamide gel strength. Ponceau stain for total protein shown as loading control. **(G)**, Proximity ligation assay (PLA) in MM134 for ER:MDC1 (left) or ER:FOXA1 (right, as control). **(H)**, PLA for ER:MDC1 as in (G). Values in G-H represent mean foci/cell +/- SD; complete quantification for (G-H) and statistical comparisons are shown in Supplemental Figure 3.

We confirmed that MDC1 (mediator of DNA damage checkpoint 1) interacts with ER by co-IP, using a different ER antibody (D8H8) than used for RIME (sc-543); ER:MDC1 interaction was observed in ILC cell lines MM134 and 44PE but not in IDC cell line MCF7 (**Figure 2C**). ER:MDC1 interaction was also confirmed via the reciprocal IP of MDC1, and we found that the ER:MDC1 interaction is not DNA-dependent as co-IP was not ablated by treatment with benzonase or ethidium bromide (**Figure 2D**). Notably, ER:MDC1 interaction was also observed in ER RIME from Martin et al using 44PE and an anti-estrogen resistant 44PE variant with mutant ER (Y537S), as MDC1 was among the top 10 proteins identified by peptide count in both models (**Figure 2E**) [30]. We further examined ER:MDC1 interaction in a panel of ER+ cell lines, and first confirmed that MDC1 is expressed across ILC cell lines (MM134, MM330, 44PE, CAMA1) and IDC cell lines (MCF7, T47D, HCC1428), and that MDC1 levels are suppressed by siRNA (**Figure 2F**; doublet v single band at 250kD is due to gel strength, data not shown). Proximity ligation assay (PLA) confirmed ER:MDC1 interaction in MM134, consistent with RIME and co-IP data; PLA signal was ablated by siMDC1 or siESR1 which confirms assay/antibody specificity (**Figure 2G**). In the cell line panel, ER:MDC1 PLA signal was detected in all ILC models and suppressed by target knockdown, consistent with ER:MDC1 proximity/interaction. In IDC models, lower PLA signal was observed but was not suppressed by siRNA, consistent with non-specific signal (**Figure 2H**; quantification and statistics in **Supplemental Figure 3**). Taken together with RIME and co-IP data, this supports that ER:MDC1 interaction is specific to or enriched in ILC cells.

Paralleling these interaction data, MDC1 knockdown suppressed proliferation in four ILC cell lines (MM134, 44PE, CAMA1, MDA MB 330/MM330; **Figure 3A-B**). In IDC cell lines, MDC1 knockdown suppressed proliferation in MCF7, but not in HCC1428 or T47D (**Figure 3A**). We confirmed that individual constructs from the siMDC1 pool utilized suppress ILC cell proliferation (**Figure 3B**). Importantly, suppression of proliferation by MDC1 knockdown is incongruous with canonical functions of MDC1 in DNA damage response, where MDC1 knockdown or loss ablates cell cycle checkpoints and allows unchecked proliferation [31, 32]. However, consistent with the effects of cell proliferation, siMDC1 halted cell cycle progression in MM134 (ILC) cells causing a G2/M arrest (**Figure 3C**). This G2/M arrest upon MDC1 knockdown contrasts G1/S arrest upon ER knockdown, suggesting MDC1 plays a discrete role in ER-mediated cell proliferation. siMDC1 did not impact cell cycle distribution in MCF7 cells, suggesting that the decrease in proliferation in MCF7 after MDC1 knockdown is related to loss of DNA repair capacity (siMDC1 also did not impact cell cycle in HCC1428, not shown). Taken together, our data suggest MDC1 is uniquely critical for ER function in ILC cells, i.e. in the context of ER:MDC1 interaction.

**Figure 3.**
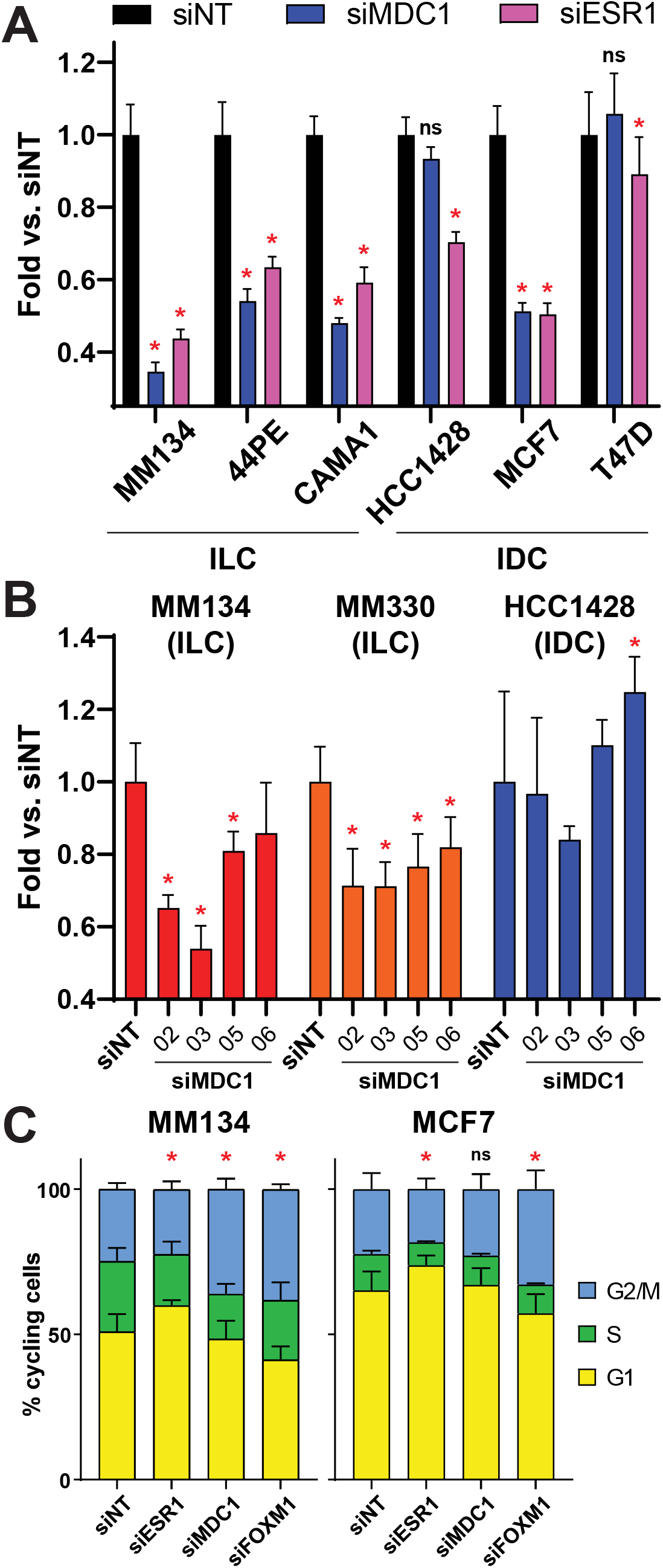
MDC1 is required for ER-mediated proliferation in ILC cells. **(A-B)**, Cell proliferation was assessed by dsDNA quantification 6d after siRNA transfection. *, p<0.05, ns, not significant vs siNT; ANOVA with Dunnett’s multiple correction. Bars represent mean of 5-6 biological replicates +/- SD. **(B)**, Individual siRNA constructs from the siMDC1 pool were transfected. (C), Cells were fixed for cell cycle analyses 72hr after siRNA transfection. *, p<0.05, ns, not significant vs siNT; ANOVA. Bars represent mean of 3-4 biological replicates +/- SEM.

### MDC1 is required for regulation of the ER transcriptome in ILC cells

MDC1 canonically mediates DNA damage response but has also implicated as a transcriptional regulator (see Discussion), consistent with a putative role as an ER co-regulator in ILC. We found MDC1 knockdown ablated regulation of ER target genes in MM134 cells (**Supplemental Figure 4A**), blocking E2-driven induction of *WNT4, IGFBP4, PDZK1*, and *TFCP2L1*, and repression of *PDE4B*. This effect of siMDC1 was not universal, as *TFF1* induction in MM134 was unaffected. No effect was observed on these ER target genes in IDC cell line HCC1428. To define how the ER transcriptome requires MDC1, we performed RNA-sequencing (RNAseq) after MDC1 knockdown in ILC models MM134 and MM330, and IDC model HCC1428 (schema in **Figure 4A**). *ESR1* knockdown (siESR1) was included to define ER target genes, and *FOXA1* knockdown (siFOXA1) was included to compare the role of MDC1 vs FOXA1.

**Figure 4.**
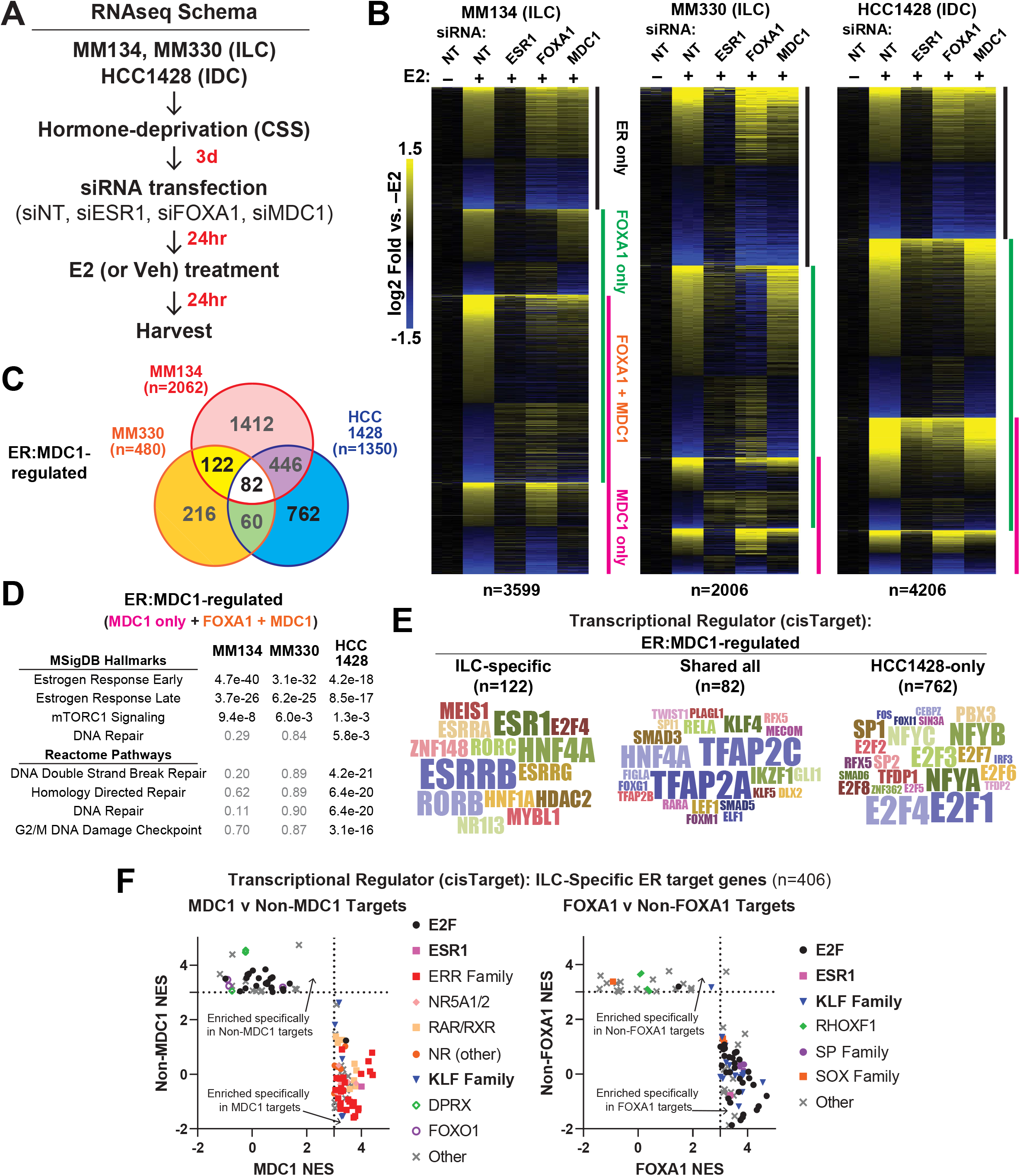
MDC1 mediates ER control of target genes associated with hormone response elements in ILC cells. **(A)**, Schema for RNA-sequencing experiment; samples were generated in biological triplicate. **(B)**, ER target genes in each cell line were identified by E2 regulation (siNT vs siNT+E2; q<0.0001) and reversal by siESR1 (siNT+E2 vs siESR1+E2, q<0.0001). For MM330, a less stringent ligand reg. cutoff was used (q<0.01) due to ligand-independent activity of ER in these cells. MDC1 and FOXA1 targets defined by the E2 effect reversed by siRNA, q<0.0001. **(C)**, Overlap of ER:MDC1 target genes across cell lines. **(D)**, ER:MDC1 target genes from (C) were subject to over-representation analyses (ORA) against MSigDB genesets (H, C2/CP, and C6 at top; C2/Reactome at bottom). **(E)**, Genes from (C) were applied to transcriptional regulator analyses (i.e. transcription factors motifs within 20kb of transcription start sites). Word clouds represent frequency of high confidence factors identified with NES≥3. Single instances of motifs are omitted from the clouds but are listed in Supplemental File 6. **(F)**, ILC-specific ER target genes were defined as MDC1- and/or FOXA1-targets as in (B) and analyzed as in (E).

We found MDC1 is required for a large proportion of the ER-driven transcriptome (**Figure 4B, Supplemental File 4**). In MM134 cells, among n=3599 ER target genes, 57.3% were dysregulated upon MDC1 knockdown, and a similar proportion were dysregulated by FOXA1 knockdown (56.1%). This included genes dysregulated by either MDC1 or FOXA1 knockdown (n=1387, 38.5%), but 18.8% (n=675) and 17.6% (n=633) of ER target genes specifically required MDC1 or FOXA1, respectively. As expected, FOXA1 knockdown also dysregulated a majority of ER target genes in MM330 and HCC1428 (54.0% and 59.9%, respectively), while MDC1 played a smaller role in the ER transcriptome in these cells compared to MM134 (23.9% and 32.1% of ER target genes in MM330 and HCC1428, respectively). siMDC1 dysregulated a large proportion of the ER transcriptome in both ILC and IDC cells, despite disparate effects on cell proliferation, but only a subset of ER:MDC1 target genes overlapped across models (**Figure 4C, Supplemental File 4**). Based on this, we examined ER-driven regulatory pathways differentially effected by MDC1 knockdown in each cell line (**Figure 4D, Supplemental File 5**). When considering all MDC1-regulated genes, the MsigDB Hallmark signatures [33] for estrogen response were enriched in all 3 models, along with mTORC1 signaling (a downstream component of ER signaling). However, Hallmark and Reactome DNA repair pathways were enriched specifically in HCC1428 (**Figure 4D**). Differential enrichment of DNA repair pathways in the ILC v IDC models suggests the role of MDC1 in ER function in IDC is related to DNA damage response [34], but in ILC cells MDC1 is potentially acting instead as an ER transcriptional co-regulator.

Similarly, we examined regulatory sequences in the promoters of ER:MDC1 target genes by querying identified target genes (from groups in **Figure 4C**) against the cisTarget database [35]. Notably, ILC-specific ER:MDC1 target genes were heavily enriched for estrogen-response-elements and half-elements (ESR1 sites and ESRR sites, respectively) and other nuclear receptor motifs (ROR, NR sites) (**Figure 4E, Supplemental File 6**). ER:MDC1 targets shared in all three cell lines were instead predominantly enriched for AP2 motifs (TFAP2A/TFAP2C), while ER:MDC1 targets unique to the IDC cell line HCC1428 were enriched for E2F motifs. Similar enrichments were not observed in individual cell lines (**Supplemental Figure 4B**) or in other overlapping pairs. The enrichment for ER sites in ER:MDC1 targets in ILC cells supports a unique role of MDC1 as an ER co-regulator in ILC cells.

Since MDC1 and FOXA1 regulate some distinct components of the ER transcriptome in ILC cells, we examined whether MDC1 versus FOXA1 target genes had distinct regulatory features in their promoter regions. We identified ER target genes for which siRNAs caused complete suppression of ER regulation (i.e. reversed expression to the estrogen-deprived baseline; **Supplemental Figure 4C-D**). From these, we identified ER target genes specific to the ILC cell lines (n=406), defined whether these genes were regulated by MDC1 and/or FOXA1, and then examined regulatory site enrichments against the cisTarget database as above (**Figure 4F, Supplemental File 7**). Paralleling our above analyses, MDC1 target genes were heavily enriched for nuclear receptor motifs, including ESR1 (ER) and estrogen-related receptors (ERR); non-MDC1 target genes were instead enriched for E2F-family motifs. Conversely, FOXA1 targets were enriched for E2F-family targets. These observations support that MDC1 plays a unique role in genes controlled by hormone-response-elements, and suggests MDC1 plays a distinct role from FOXA1 in ER-mediated gene regulation in ILC cells.

### MDC1 protein is differentially expressed in ER+ ILC tumors

Studies of MDC1 protein in human tumors suggest MDC1 is over-expressed in some tumor types to suppress apoptosis, but is more often decreased or lost in tumors relative to normal tissues, likely promoting genomic instability (recently reviewed [36]). In breast cancer, limited analyses to date have suggested MDC1 protein is decreased or lost in 30-70% of tumors, and that MDC1 loss is associated with advanced stage and poor outcomes [37–39], but associations with ER status and histology are unknown. To examine this further, we first assessed *MDC1* mRNA levels in the Cancer Genome Atlas (TCGA) and METABRIC cohorts via cBioPortal [40]. In both TCGA and METABRIC, *MDC1* expression was highest in ER-/HER2-IDC tumors compared to other IDC and to ER+/HER2-ILC (**Figure 5A**). Considering ER status only, *MDC1* expression was higher in ER-IDC vs either ER+ IDC or ER+ ILC in both datasets (FDR q<0.05); PR status was not associated with differences in *MDC1* expression in ER+ tumors (not shown). We did not identify significant associations between *MDC1* mRNA levels and outcome in either cohort (not shown). We next assessed MDC1 protein levels by IHC staining using commercial breast cancer tissue microarrays (TMAs). We investigated two antibodies for IHC staining, and found that Abcam ab11171 (Abcam) showed specific nuclear staining in MM134 FFPE cell pellets and normal breast tissue, and siMDC1 decreased staining intensity in MM134 cell pellets (**Supplemental Figure 5A**). In breast tumor tissues, MDC1 IHC showed a broad range of expression levels across IDC and ILC tumors (**Figure 5B**; **Supplemental File 8**). ANOVA analyses did not detect statistically significant differences between ILC/IDC based on ER status. However, we noted that low/negative MDC1 by IHC was rare in ER+ ILC tumors (chi-test p = 0.027), which had MDC1 levels more consistent with normal and non-malignant breast tissues (**Figure 5C, Supplemental Figure 5B**). The extent of MDC1 protein loss by IHC was not mirrored in mRNA data, suggesting that MDC1 may be post-transcriptionally regulated in breast tumors cells (consistent with MDC1 protein regulation in DDR [27, 28, 41]). While MDC1 protein loss or downregulation may be a mechanism by which tumor cells facilitate increased genomic instability, ER+ ILC may uniquely maintain MDC1 protein based on its critical role in ER function.

**Figure 5.**
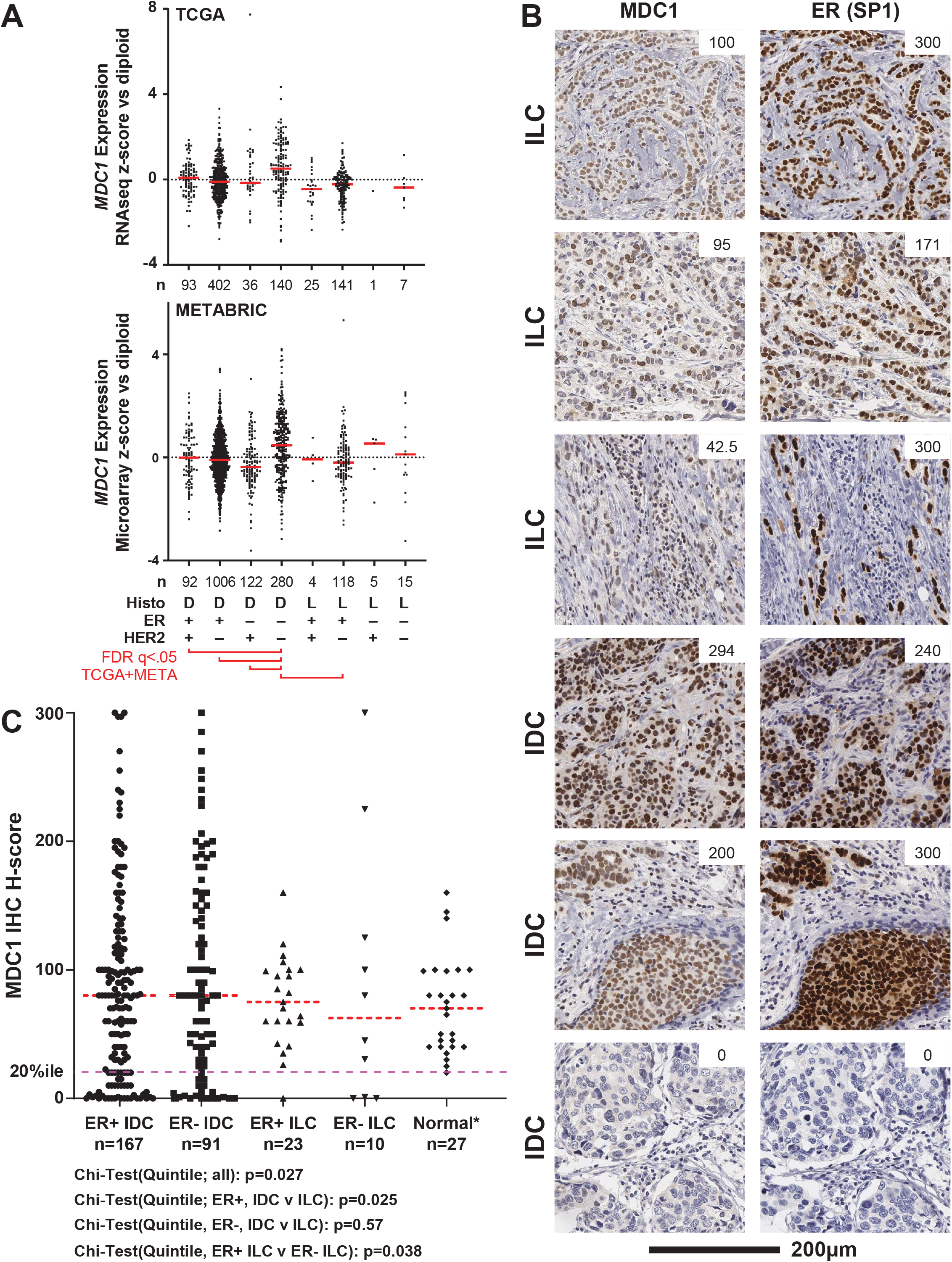
Loss of MDC1 protein is uncommon in ER+ ILC tumors. **(A)**, *MDC1* mRNA levels in breast tumors from TCGA (Firehose legacy dataset) and METABRIC cohorts. Red line = median expression. **(B)**, Representative IHC images with H-scores from breast cancer TMAs. **(C)**, MDC1 H-scores from full TMA cohort. Normal tissues include healthy/unaffected mammary glands, hyperplasia, and fibroadenoma samples (see Supplemental File 8). Dashed red line = median H-score. Pink dash line = lower quintile for full cohort.

## DISCUSSION

ILC are nearly all ER+ and have tumor biomarkers consistent with the Luminal A molecular subtype, but these tumors have a unique relationship with estrogen and anti-estrogens relative to other breast cancers. For example, hormone replacement therapy using estrogen alone is sufficient to increase the risk of ILC, whereas IDC risk is restricted to combined estrogen plus progestin therapy [1]. Despite being exquisitely estrogen-dependent, ILC may be resistant to anti-estrogens, as suggested by limited response to tamoxifen (clinically [10, 11] and in lab models [12, 13, 42, 43]) and by the poor long-term outcomes for patients with ILC [5–8]. Based on these data, we hypothesized that ER function is unique in ILC cells and examined novel and/or ILC-specific ER co-regulator proteins as a driver of ER functions in ILC cells. Using the co-IP/MS method RIME, coupled with an siRNA screen against ER-driven proliferation, we identified the DNA damage response protein MDC1 as an ER-associated protein in ILC cells. MDC1 is required for ER-driven proliferation in ILC cells, with MDC1 knockdown causing cell cycle arrest, whereas in the context of DDR function MDC1 knockdown or loss abrogates cell cycle checkpoints. Similarly, RNAseq analyses showed MDC1 is required for a major subset of the ER-regulated transcriptome in ILC cells, and is associated with DNA repair signatures only in IDC cells, suggesting that MDC1 has ER co-regulator functions independent of its canonical role in DDR. Further, by IHC we observed that loss of MDC1 protein is uncommon specifically in ER+ ILC, consistent with a critical role for MDC1 in supporting ER function in ILC cells.

Beyond the canonical roles of MDC1 in DDR, reports in the literature implicate MDC1 as having coregulator activity. In GAL4 fusion assays, MDC1 (a.k.a. NFBD1 or KIAA0170) residues 508-995 enhanced reporter activity in COS cells [44]. Other studies link MDC1 to epigenomic remodeling proteins, including the SWI/SNF complex [45] and p300 histone acetyltransferase [46, 47]. Interaction with epigenomic enzymes may suggest MDC1 acts to scaffold proteins on ER to mediate gene regulation (MDC1 has no known enzymatic activity), similar to the NCOA family of co-regulators [48]. However, the canonical roles of MDC1 in DDR could also impact ER-driven gene regulation, since ER may regulate genes by causing DNA damage to induce DDR-driven recruitment of chromatin remodelers [49]. In this context, MDC1 has been previously reported to interact with ER and AR (in MCF7 and CWR22Rv1 prostate cancer cells, respectively), but in these settings MDC1 acted as a tumor suppressor [36, 39, 50]. In breast and prostate cancer lines, MDC1 shRNA knockdown enhanced proliferation, migration, and invasion, consistent with abrogation of DDR cell cycle checkpoints. MDC1 is also a tumor suppressor in tumorigenesis models, such as in *Mdc1*-knockout mice which show increased tumorigenesis due to increased genomic instability [51–54]. MDC1 also functions as a tumor suppressor in DDR in ER-negative breast cancer, where MDC1 links ID4 to the BRCA1 DDR network [55]. However, our data contrast the tumor suppressor roles of MDC1 in DDR, and instead suggest an oncogenic role in mediating ER function and ER-mediated proliferation. Similarly, we found *MDC1* siRNA did not block AR-driven (ER-independent) growth in MM134, supporting a specific role in ER function in ILC cells. Taken together, MDC1 can contribute to nuclear receptor function via DDR and as a tumor suppressor in some settings, but in ILC, MDC1 has unique and potentially oncogenic functions as an ER co-regulator.

Understanding how MDC1 has ER co-regulator activity specifically in ILC cells is an important future direction, and may be facilitated by differential post-transcriptional modification of MDC1 in the context of ILC. MDC1 has extensive post-translational modification (PTM) including SUMOylation [56], methylation [57], phosphorylation [31], and ubiquitination [27, 28], and MDC1 cleavage can generate MDC1 unable to engage in DDR [41]. Future studies will examine differential MDC1 PTM in ILC versus IDC contexts. Differential MDC1 PTM as a mechanism to facilitate ER interaction may parallel our recent findings that ER protein is differentially regulated in ILC cells, showing reduced ligand-driven ubiquitination and degradation versus ER in IDC cells [16]. Defining how MDC1 and ER interact together on chromatin is also critical to understanding the ILC-specific co-regulator functions of MDC1. However, MDC1 chromatin binding in DDR [44, 58] creates technical issues for ChIP studies. We find different MDC1 antibodies have varying ability to co-IP ER versus known DDR partners (not shown), and the latter may confound the interpretation of ChIP data of ER:MDC1. Future studies will better define the ER:MDC1 interaction, to both understand the ILC-specificity of the interaction and facilitate genomic studies of ER:MDC1 function.

The putative oncogenic functions of MDC1 as an ER co-regulator in ILC raises questions as to whether MDC1 maintains its tumor suppressor functions in DDR in ILC cells. In our RNAseq, ER:MDC1 target genes in ILC cells were not enriched for DDR-related signatures or E2F promoter motifs (unlike our data from IDC cells), which may suggest a disconnect between MDC1 and DDR in ILC cells. Similarly, TCGA data suggest DDR may be dysfunctional in some ILC tumors. TCGA identified a ‘Proliferative’ ILC subtype with the poorest disease-free and overall survival (2-fold increased recurrence)[2]. Proliferative ILC has an elevated DDR signature (by reverse phase protein array, RPPA) vs other ILC subtypes or vs Luminal A IDC, defined by high DDR protein levels [59], but the functional implications of this elevated DDR signature has not been explored. Clinically, ILC have the poorest response to chemotherapy among breast cancer subtypes, and patients with ILC receive little if any survival benefit from chemotherapy [60–63]. Further mechanistic studies are needed to understand chemo-response and -resistance, DDR function, and the potential crosstalk between ER, MDC1, and DDR in ILC cells.

Similarly, our data on the critical role of MDC1 in tamoxifen-mediated ER function, and identification of MDC1 as a partner of mutant ER in anti-estrogen resistant/ER mutant 44PE cells, suggest that MDC1 plays a central role in ER-dependent anti-estrogen resistance. Defining the role of ER:MDC1 interaction in clinical anti-estrogen response and patient outcomes may require functional assessment of MDC1 in tumors tissues; our data suggest *MDC1* mRNA and MDC1 protein are not strongly correlated, and protein levels by IHC alone likely cannot confirm ER:MDC1 interaction or MDC1 co-regulator activity. Additional data on estrogen-mediated gene regulation in ILC tumors can facilitate identifying ER:MDC1-regulated genes *in vivo*, such as gene expression data from an ongoing trial of neoadjuvant anti-estrogen therapy for patients with ILC [NCT02206984].

Increasing evidence from clinical, epidemiological, and laboratory studies suggest that ILC is a biologically distinct subset of breast cancer, and further that ILC represents a unique context for ER activity. Our understanding of ER and anti-estrogens in breast cancer is almost exclusively based on IDC biology (e.g. data from MCF-7), but work from our lab and others supports that ILC should be considered an independent tissue type, not unlike study of ER in the endometrium, bone, CNS, or other tissues. As seen in other tissues, distinct co-regulator cohorts may mediate tissue type-specific ER functions in ILC, and we identified MDC1 as a unique ER co-regulator in ILC cells that is critical in estrogen response and anti-estrogen resistance phenotypes. Further defining MDC1 functions will provide mechanistic basis for the unique ER activity in ILC, thus potentially underlying the distinct biology of ILC. MDC1 may be a therapeutic target for a patient population in critical need of precision treatments.

## MATERIALS AND METHODS

### Cell culture and reagents

MDA MB 134VI (MM134; ATCC, Manassas, VA, USA) and SUM44PE (Asterand/BioIVT, Westbury, NY, USA) were maintained as described [12]. MDA MB 330 (MM330; ATCC) and HCC1428 (ATCC) were maintained in DMEM/F12 (Corning #10092CV) + 10% fetal bovine serum (FBS; Nucleus Biologics #FBS1824). MCF7 and CAMA-1 lines were generous gifts from the Rae Lab at the University of Michigan, and were maintained in DMEM/F12 + 10% FBS. T47D were a generous gift from the Sartorius Lab at the University of Colorado, and were maintained in MEM (Corning #10010) + 5% FBS + 1x Non-essential amino acids + 1nM sodium pyruvate + 1nM insulin. BCK4 were a generous gift from the Jacobsen Lab at the University of Colorado and were maintained as described [64]. Hormone-deprivation was performed as described [65] with phenol red-free reagents in IMEM (ThermoFisher #A10488) supplemented with 10% charcoal-stripped fetal bovine serum (CSS; prepared as described [65] with the same FBS as above). All lines were incubated at 37°C in 5% CO_2_. All cell lines were regularly confirmed to be mycoplasma negative and were authenticated by STR profiling at the University of Colorado Anschutz Tissue Culture Core. Estradiol (E2), 4-hydroxytamoxifen (4OHT), and testosterone (TS) were obtained from Sigma; ICI 182780 (fulvestrant; fulv) and synthetic androgen Cl-4AS-1 were obtained from Tocris Biosciences. Small molecules were dissolved in ethanol, and vehicle treatments are using 0.1% EtOH.

### siRNA Knockdown Protocol

siRNA constructs were reverse transfected using RNAiMAX (ThermoFisher) according to manufacturer instructions. Constructs are siGENOME SMARTpool siRNAs (GE Healthcare Dharmacon, Lafayette, CO, USA) containing 4x siRNA targeting constructs, or individual siRNA constructs: Non-targeting pool #2 (D-001206-14-05), Human *ESR1* pool (M-003401-04-0010), Human *MDC1* pool (M-003506-04-0005), Human *FOXA1* pool (M-010319-01-0005), and Human *MDC1* individual constructs (D-003506-02-0002, D-003506-03-0002, D-003506-05-0002, D-003506-06-0002).

### Rapid immunoprecipitation mass spectrometry of endogenous proteins (RIME)

RIME was performed with Active Motif (Carlsbad, CA), and samples were prepared according to the commercial protocol (**Supplemental File 9**). Briefly, MM134, 44PE, or BCK4 cells were plated in standard conditions containing FBS (above) in three 15cm plates; 1 plate each was used for Vehicle treatment (ER IP), 4OHT treatment (ER IP), or Vehicle treatment for IgG IP control. Cells were treated with Vehicle (0.1% EtOH) or 4OHT (1µM) for 24hr prior to harvest. At the time of harvest, cells (MM134: ∼9×10^7^/plate; 44PE: ∼3×10^7^/plate; BCK4: ∼2.5×10^7^/plate) were fixed in 11% formaldehyde solution for 8min at room temperature with gentle rocking. Fixation was quenched with 1/20 volume of 2.5M glycine, and cells were collected by scraping. After centrifugation, pellets were washed twice in 0.5% Igepal CA-630 + 1mM PMSF in PBS, then pelleted and snap frozen. Nuclear isolation, immunoprecipitation, and mass spectrometry (in technical duplicates) were performed by Active Motif. ER IP used antibody sc-543 (Santa Cruz Biotechnology; Santa Cruz, CA; antibody is discontinued) and rabbit IgG. MS was performed in technical duplicate for each sample (single biological replicate per condition). Quality control reports from the mass spectrometry analyses are provided in **Supplemental File 10**.

Identified peptides were filtered based on having <20% of peptides for a given protein in IgG samples relative to all samples, and the identified protein being present in <40% of studies in the CRAPome database [66]. Common tubulin and ribosomal protein contaminants were also excluded. ER RIME data from IDC cells in Mohammed et al [26] used the union of peptides identified across biological replicates.

### siRNA screen

siRNA SMARTpools containing 4 siRNA constructs per pool were ordered as a pre-plated custom libraries in 96-well format (Dharmacon / Horizon Discovery; Layfayette, CO; sequences listed in **Supplemental File 3**). MM134 cells were hormone-deprived as described above prior to reverse transfection with 10nM siRNA (as above). 24hr post-transfection, cells were treated with Vehicle (0.01% EtOH), 100pM E2, 100nM 4OHT, or 10nM Cl-4AS-1 (all in biological triplicate) and allowed to grow for 6 days prior to assessing proliferation by dsDNA quantification (see below). Raw data from dsDNA quantification and associated statistical analyses and target filtering are provided in **Supplemental File 3**. Data analyses were performed in Graphpad Prism v7.0.

### Cell proliferation

Total double-stranded DNA was measured as a surrogate for total cell number. dsDNA was quantified by hypotonic lysis of cells in ultra-pure H_2_O, followed by addition of Hoechst 33258 (ThermoFisher Scientific, #62249) at 1µg/mL in Tris-NaCl buffer (10mM Tris, 2M NaCl; pH 7.4) at equivalent volume to lysate. Fluorescence (360nm ex / 460nm em) was measured on a Bio-Tek Synergy 2 microplate reader.

### Immunoblotting

Cells were seeded in 12- or 24-well plates for ∼80% confluence according to cell size (MM134 and SUM44PE: 800k/well or 400k/well; MM330: 640k/well or 320k/well; CAMA-1: 400k/well or 200k/well; MCF7 and T47D: 320k/well or 160k/well; HCC1428: 240k/well or 120k/well) and treated with 10nM siRNA, 100pM E2, and other treatments as indicated. Cell lysates were harvested after treatments in RPPA buffer (50mM HEOES pH 7.4, 150mM NaCl, 1.5mM MgCl_2_, 1mM EGTA, 10% Glycerol. 1% Triton X-100, dH_2_O) with phosphatase inhibitors (ThermoFisher #78420). BioRad Mini-PROTEAN TGX precast gels were used for electrophoresis and transferred to PVDF membranes. Membranes were blocked in either 5% milk or 5% BSA in Tris-buffered saline with 0.5% Tween 20, depending on primary antibody used. Primary antibody probing was performed according to manufacturer recommended concentrations, and species-matched secondary antibodies were diluted in TBS + 0.05% Tween-20. Membranes were imaged on LiCor C-DiGit blot scanner. Primary antibodies for immunoblots were: ER, Leica 6F11 (Leica Biosystems #PA009); MDC1, Sigma M2444.

### Co-immunoprecipitation (Co-IP)

Cells were seeded in 10cm plates for ∼80% confluency as described above. All centrifugation steps below were at 4°C. Cultures were rinsed and then harvested by scraping in HBSS+1mM EDTA, then centrifuged for 5’ at 1000xg. Pellets were rinsed in 1mL of PBS and centrifuged for 3’ at 1000x*g*. The cell pellet was re-suspended in 3 packed cell volumes of Hypotonic Buffer A (10mM HEPES-KOH pH 7.6; 1.5mM MgCl_2_, 10mM KCl, 1x protease inhibitors, and 1mM DTT). Cells were incubated on ice for 3’ prior to Dounce homogenization; nuclei were then centrifuged for 15’ at 18000x*g*. The supernatant was reserved as the ‘cytoplasmic fraction’, and nuclei were re-suspended in two packed nuclear volumes (PNV) of Hypertonic Buffer B (20mM HEPES-KOH pH 7.6, 25% glycerol, 1.5mM MgCl_2_, 0.6M KCl, 0.2mM EDTA, 1x protease inhibitors, and 1mM DTT), homogenized by pipetting, and incubated at 4°C for 45’ with rotation. Resulting lysate was centrifuged for 10’ at 18000x*g* to collect the supernatant. Supernatant was diluted with 3 PNVs of Hypertonic Diluent Buffer C (20mM HEPES-KOH pH7.6, 1.5mM MgCl_2_, 0.2mM EDTA, 0.67% NP-40 (IGEPAL), 1x protease inhibitors, and 1mM DTT), then insoluble proteins were removed by centrifuging for 5’ at 18000x*g*. The resulting supernatant (nuclear extract, NE) was utilized for immunoprecipitation.

NE was combined with 10ug of antibody for immunoprecipitation. Jackson ImmunoResearch Chromapure Rabbit or Mouse served as IgG controls. α-MDC1 (Sigma, M2444) or α-ER (Cell Signaling, clone D8H8, Cat # 8644) were used for reciprocal co-IP. NE was incubated with antibody overnight at 4°C with rotation. Protein A/G magnetic beads (Thermo Scientific / Pierce, Cat # 88803; 25ul/10ug antibody) were prepared by washing in Buffer D (20mM HEPES-KOH, pH7.6, 1.5mM MgCL_2_, 150mM KCl, 0.2mM EDTA, and 0.67% NP-40 (IGEPAL)) and separation by magnetic stand. Beads were added to IP reactions and incubated for 4 hours at 4^0^C with rotation. Beads were washed 5 times with 1mL of Buffer D. Protein complexes were eluted by re-suspending beads in 100ul of Buffer D supplemented with 6x Laemmli buffer and allowed to incubate for 10’ at room temperature. Beads were then removed via magnetic stand, and cleared lysates were denatured by heating at 95°C for 10’, then assessed by immunoblot as above.

For benzonase and ethidium bromide (EtBr) co-IP experiments, Buffer B was supplemented with 3uL of benzonase (Millipore Sigma; ≥250units/uL, Cat #E1014) or 100ug/mL of Ethidium bromide (Bio Rad, Cat #1610433).

### Proximity Ligation Assay (PLA)

Cells were seeded for optimum density according to cell size (MM134 and SUM44PE at 40k/well, MM330 32k/well, CAMA-1 at 20k/well, MCF7 and T47D at 16k/well, HCC1428 at 24k/well) in 10-well chamber coverslides (Greiner Bio-One #543079) and incubated for 48 hours with indicated drug or siRNA at 37C at 5% CO_2_. Cells were washed 2x with room temperature 1x PBS and fixed in 4% paraformaldehyde (Electron Microscopy Sciences #15710) for 15min with shaking at room temperature. Cells were washed 2x with room temperature 1x PBS. Cells were permeabilized with 0.1% Triton X-100 (ThermoFisher #BP151-100) in 1x PBS for 15min at room temperature with shaking and rinsed in 1x PBS. PLA was then performed according to Sigma Aldrich Duolink PLA Fluorescence Protocol (DUO92008, DUO92002, DUO92004). Primary antibodies for proteins of interest were diluted to 1:200 (ER 6F11) or 1:2500 (Bethyl MDC1 A300-051A). DNA was stained for imaging using Duolink mounting medium with DAPI (Sigma #DUO82040). PLA foci were quantified using JQuantPlus software [67].

### Cell cycle analyses

Cells were plated in 6-well plates and transfected with siRNA as above. 72 hours post-siRNA, cells were harvested by trypsinization, then fixed in 500uL of ice cold 70% ethanol at 4°C for 30’. Cells were centrifuged at for 3’ at 18000x*g* and washed twice in 1x PBS, then stained in 500uL of FACS buffer (1x PBS, Ca^2+^/Mg^2+^ free, 1mM EDTA, 25mM HEPES pH 7.0 and 1% BSA) supplemented with 10µg of RNase A (ThermoFisher) and 10µg of Propidium Iodide solution. Cells were incubated overnight at 4°C prior to analysis on a Gallios cytometer (Beckman Coulter Life Sciences) using Kaluza 2.1 analysis software (Beckman Coulter).

### Quantitative PCR

RNA extractions were performed using the RNeasy Mini kit (Qiagen). RNA was converted to cDNA using Promega reagents: 1:1 Oligo (dT)_15_ primer (cat# C110A) + Random hexamers (cat# C118A); GoScript 5x Reaction Buffer (cat# A500D); MgCl_2_ (cat# A351H), dNTP (cat# U1511); RNasin Plus (cat# N261B); GoScript Reverse Transcriptase (cat# A501D). qPCR reactions were performed with PowerUp SYBR Green Master Mix (Life Technologies, cat #100029284) on a QuantStudio 6 Flex Real-Time PCR system. Expression data were normalized by ΔCt to *RPLP0*. Primers used are listed in **Supplemental File 11**.

### RNA sequencing and analysis

MM134, MM330, and HCC1428 cells were hormone deprived according to protocol (see above), plated for optimal confluency in 48-well plates, and transfected with 10nM siNT, siMDC1, siER, and siFOXA1 according to the method described above. 24hours after siRNA transfections, cells were treated with 100pM E2. Cells were harvested 24 hours after addition of E2, washed 3X with cold PBS, and frozen at -8°C. RNA was extracted as above. Knockdown validation via qPCR was performed before sample submission.

Library preparation and sequencing was performed at the University of Colorado Comprehensive Cancer Center Microarray and Genomics Core Facility. Libraries were generated with Illumina Stranded mRNA Prep, and were sequenced on a NovaSEQ6000 with 2×150 paired-end reads, targeting 40×10^6^ clusters / 80×10^6^ reads per sample (final mean = 39,019,637 clusters/sample). Sequencing data were analyzed with the University of Colorado Comprehensive Cancer Biostatistics and Bioinformatics Shared Resource. Illumina adapters and the first 12 base pairs of each read were trimmed using BBDuk [68] and reads <50bp post trimming were discarded. Reads were aligned and quantified using STAR (2.6.0a) [69] against the Ensembl human transcriptome (hg38.p12 genome (release 96)). Reads were normalized to counts per million (CPM) using the edgeR R package [70]. Differential expression was calculated using the limma R package and the voom() function [71]. Over-representation analysis was performed with the clusterProfiler R package [72] and gene sets from the Molecular Signatures Database [33]. Transcriptional regulators were inferred using the RcisTarget R package [73] and the cisTarget database [35]. A normalized enrichment score (NES) of 3 was used as a cutoff. Heatmaps were generated with MeV v4.9.0 [74]. Raw and processed data will be deposited in GEO upon manuscript submission.

### Immunohistochemistry

Tissue microarrays were acquired from Pantomics (Fairfield, CA): BRC1021, BRC1507, BRC15010, BRM961, BRM962. IHC was performed by the CU Anschutz Research Histology Shared Resource. Antigen retrieval used citrate buffer pH9.0 with 30’ incubation, followed by staining for ER (SP1, 1:100) or MDC1 (Abcam ab11171, 1:500; Bethyl A300-051, 1:1000). Staining was assessed by a board-certified pathologist (SM) and converted to an H-score. For a subset of tumors with duplicate cores, H-scores were averaged across the duplicates. Detailed information and scoring is provided in **Supplemental File 8**.

## SUPPLEMENTAL MATERIALS

The following supplemental materials are available for download:

- Supplemental Figures 1–5
- Supplemental File 1 (xlsx): MS spectrum counts related to Figure 1A-B
- Supplemental File 2 (xlsx): siRNA panel selections
- Supplemental File 3 (zip): siRNA screening raw data and analyses
- Supplemental File 4 (xlsx): ER target gene lists and regulation designation related to Figure 4B
- Supplemental File 5 (xlsx): Pathway analyses of ER:MDC1 target genes related to Figure 4D
- Supplemental File 6 (xlsx): Transcriptional reg analyses of MDC1 targets related to Figure 4E
- Supplemental File 7 (xlsx): Transcriptional reg analyses of MDC1vFOXA1 targets related to Figure 4F
- Supplemental File 8 (xlsx): TMA scoring and information
- Supplemental File 9 (pdf): Active Motif RIME protocol
- Supplemental File 10 (pdf): Active Motif quality control reports for ER RIME
- Supplemental File 11 (xlsx): qPCR primers used in Supplemental Figure 4

## ACKNOWLEDGEMENTS

The authors thank Dr. Mary Reyland and Angela Ohm for technical assistance in utilizing the JQuantPro software.

## DATA AVAILABILITY

RIME data (including QC data and Scaffold files provided by Active Motif) and RNA sequencing data are available upon request and will be deposited to public repositories on publication.

---

**Supplemental Figure 1.**
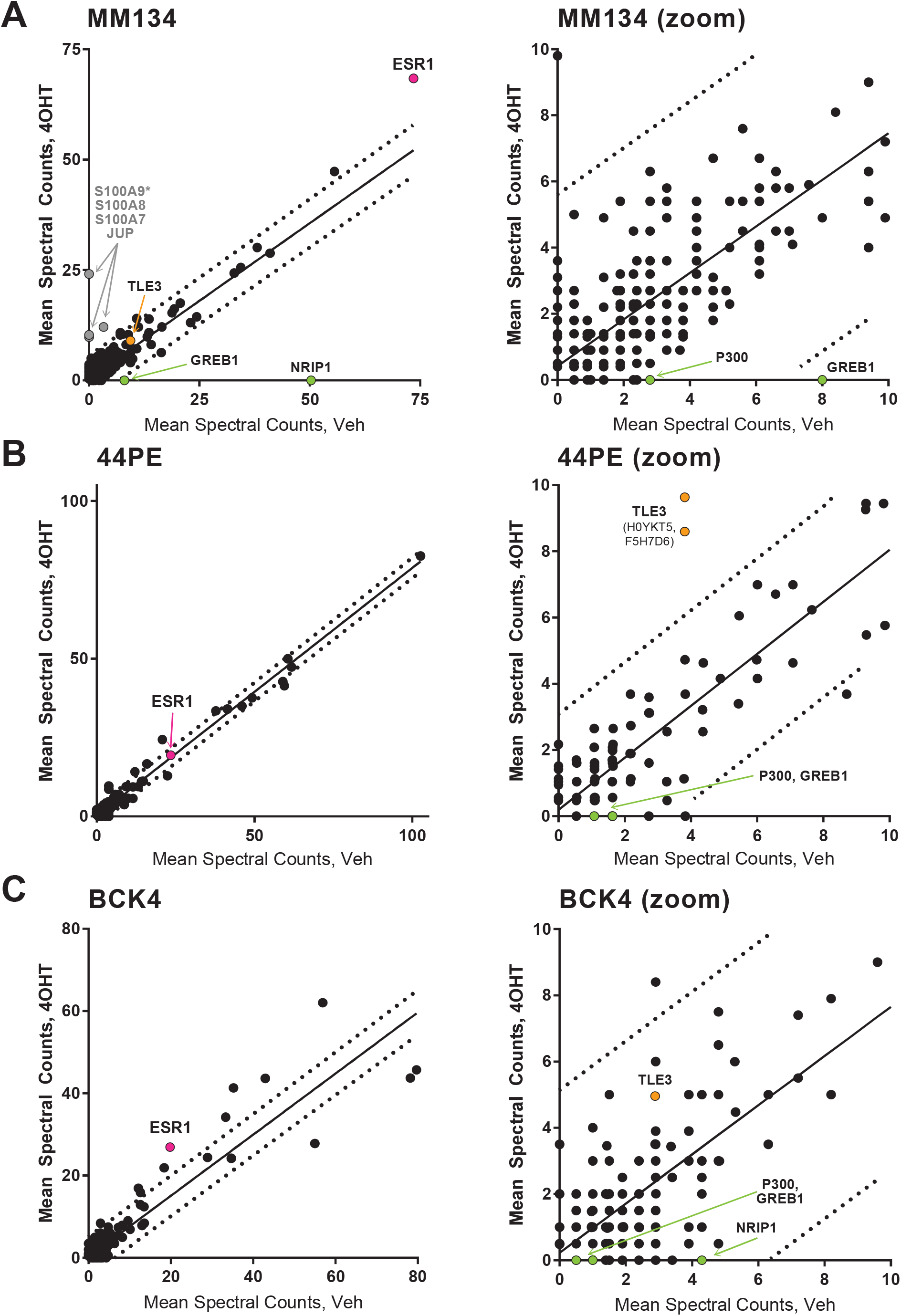
Tamoxifen treatment does not substantially alter the cohort of protein associated with ER. Points represent mean of technical replicates for proteins identified in Vehicle- vs 4OHT-treated cells for **(A)** MM134, **(B)** 44PE, or **(C)** BCK4. Lines represent linear regression and 95% prediction bands. ER and known ER co-regulators are labeled. *, proteins were identified in a single technical replicate.

**Supplemental Figure 2.**
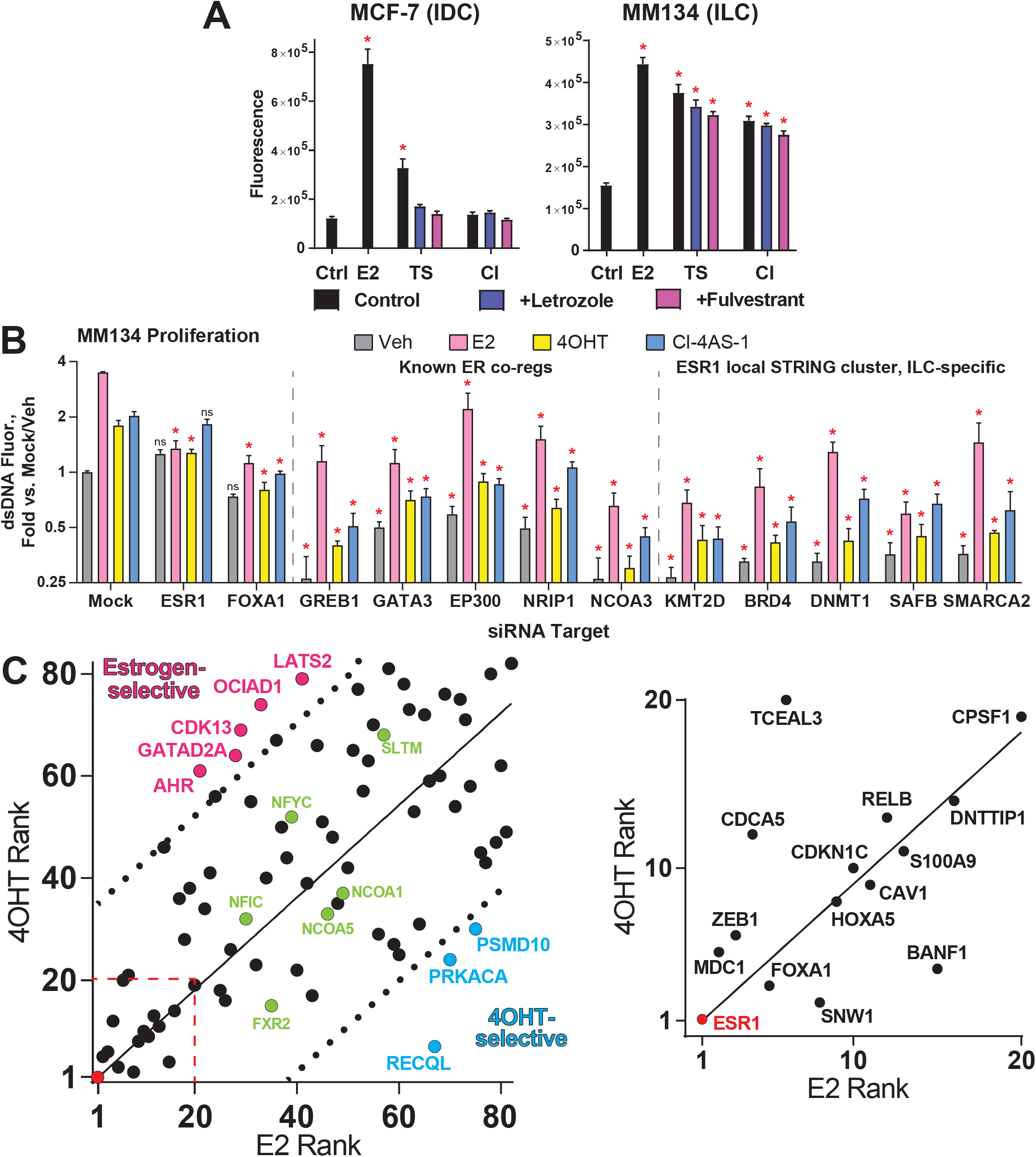
Selectivity for ER-associated proteins per ER ligand and effects on proliferation. **(A)**, Cells were hormone-deprived prior to treatment with 100pM E2, 100nM testosterone (TS) or Cl-4AS-1 (Cl; non-aromatizable synthetic androgen), +/- 1µM fulvestrant or letrozole (aromatase inhibitor). Bars = mean of 5 biological replicates +/-SD. *, p<0.05 vs Control (far left), ANOVA + Dunnett’s correction. **(B)**, Examples of known co-regulators that were filtered out by suppression of Veh/-E2 growth and AR-driven growth. Bars = mean of biological triplicate +/-SD. *, p<0.05 vs matched Mock siRNA condition, ANOVA + Dunnett’s correction. ns, not significant. ESR1 and FOXA1 shown at left as key examples passing filter criteria. **(C)**, siRNA targets that passed stringency filters (n=82, including ESR1; see Figure 2A) were ranked by their suppression of E2-versus 4OHT-induced growth. Dotted lines indicate 90% prediction bands based on linear regression (regression forced through 0,0). Dashed red box is shown as zoomed inset at right.

**Supplemental Figure 3.**
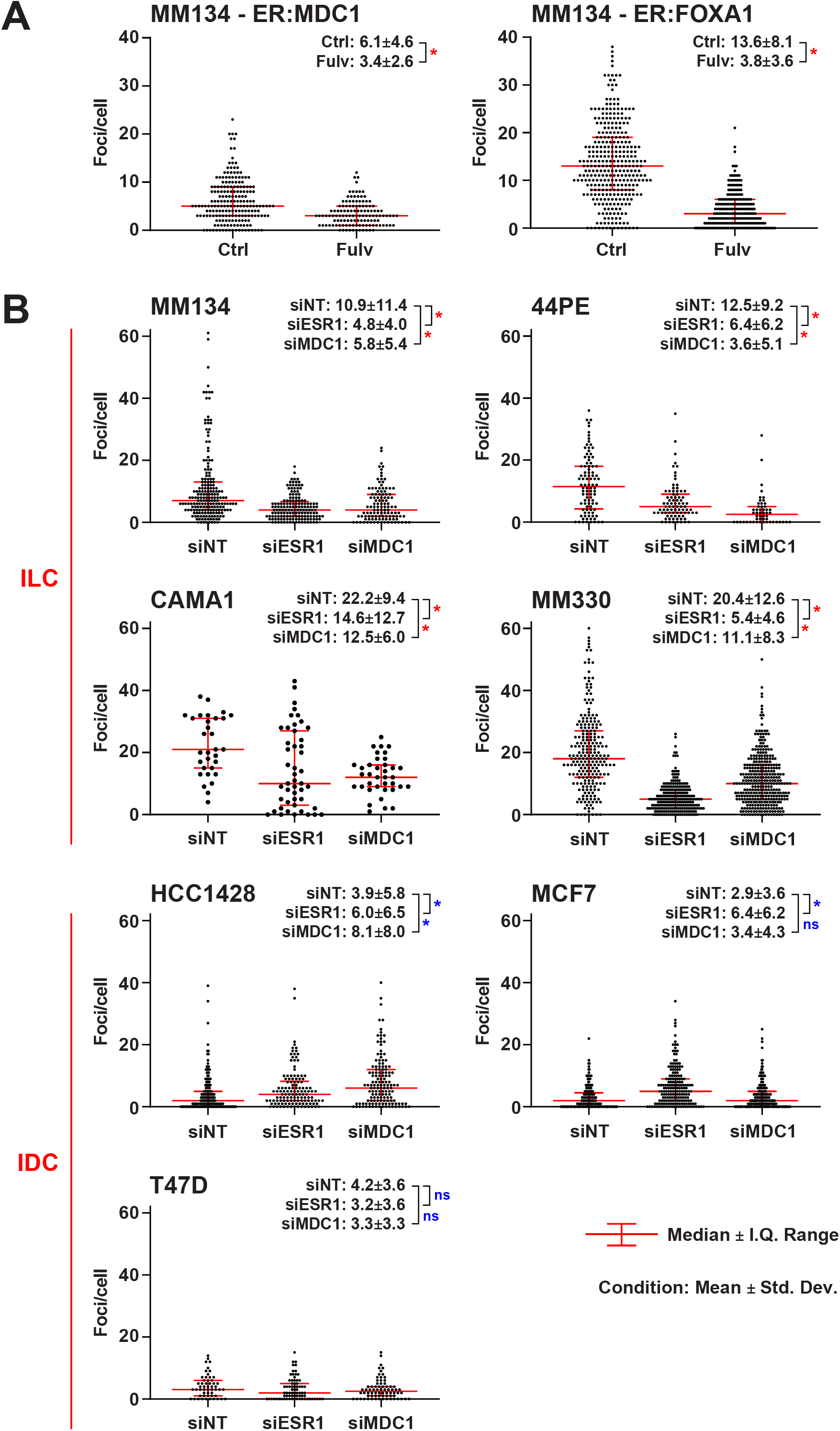
ER:MDC1 PLA foci counts are suppressed by ER or MDC1 knockdown in ILC cells. Foci were quantified using JQuantPro software, using DAPI signal as a mask for nuclei (only nuclear PLA signal was quantified). Foci parameters (e.g. signal intensity and size cutoffs) were developed using the samples in (A), then applied to samples in (B) with minor modifications per cell line. **(A-B)**, MM134 were treated with 100nM fulvestrant or siRNA as indicated prior for 48h to PLA. Points represent foci count per individual nucleus. Red lines show median foci/nucleus +/- interquartile range. P-values represent siRNA vs siNT, non-parametric ANOVA with Dunn’s multiple testing correction. **(A)**, Fulvestrant leads to ER protein degradation, acting as a control for PLA signal specificity. **(B)**, ER:MDC1 PLA quantification as in (B). Blue values note a non-significant change or a statistically significant increase in PLA foci after siRNA, inconsistent with specific ER:MDC1 PLA signal.

**Supplemental Figure 4.**
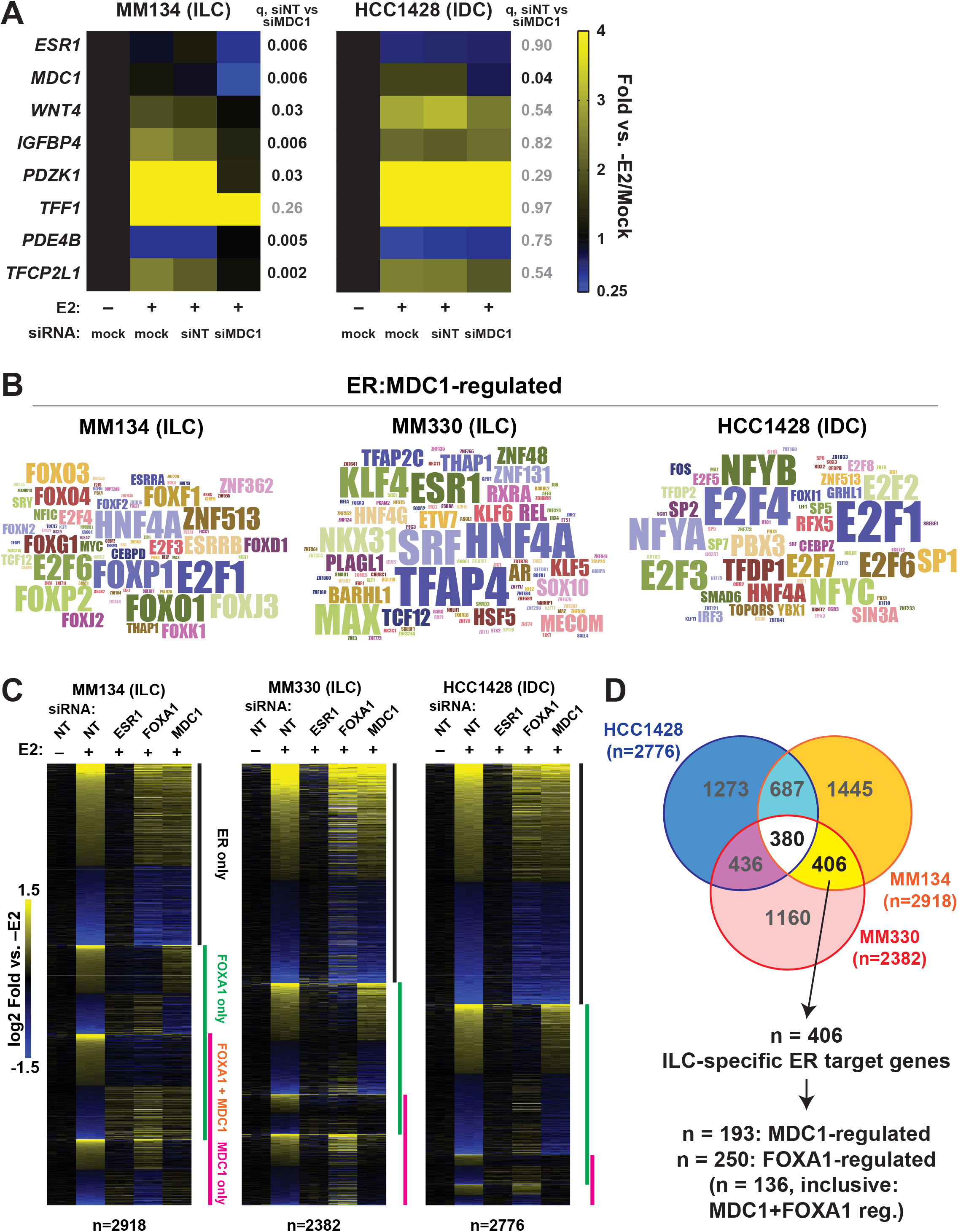
MDC1 mediates ER control of target genes associated with hormone response elements in ILC cells. **(A)**, Cells were hormone-deprived prior to siRNA transfection; 24hrs later cells were treated +/-100pM E2 for an additional 24hrs. Samples were generated in biological triplicate, mean shown in heatmap. q-value represents ANOVA with Dunnett’s multiple testing correction. **(B)**, Word clouds representing frequency of high confidence factors identified with NES≥2.5 in all ER:MDC1 targets from the indicated cell line. **(C)**, ER targets were defined by siRNA fully suppressing E2-induced expression (i.e. siRNA+E2 vs siNT, q>0.05; equivalent to -E2). **(D)**, Overlapping ER target genes from (C) identifies high-confidence ILC-specific ER target genes. MDC1 and FOXA1 targets were defined by siRNA fully suppressing E2-induced expression as in (C).

**Supplemental Figure 5.**
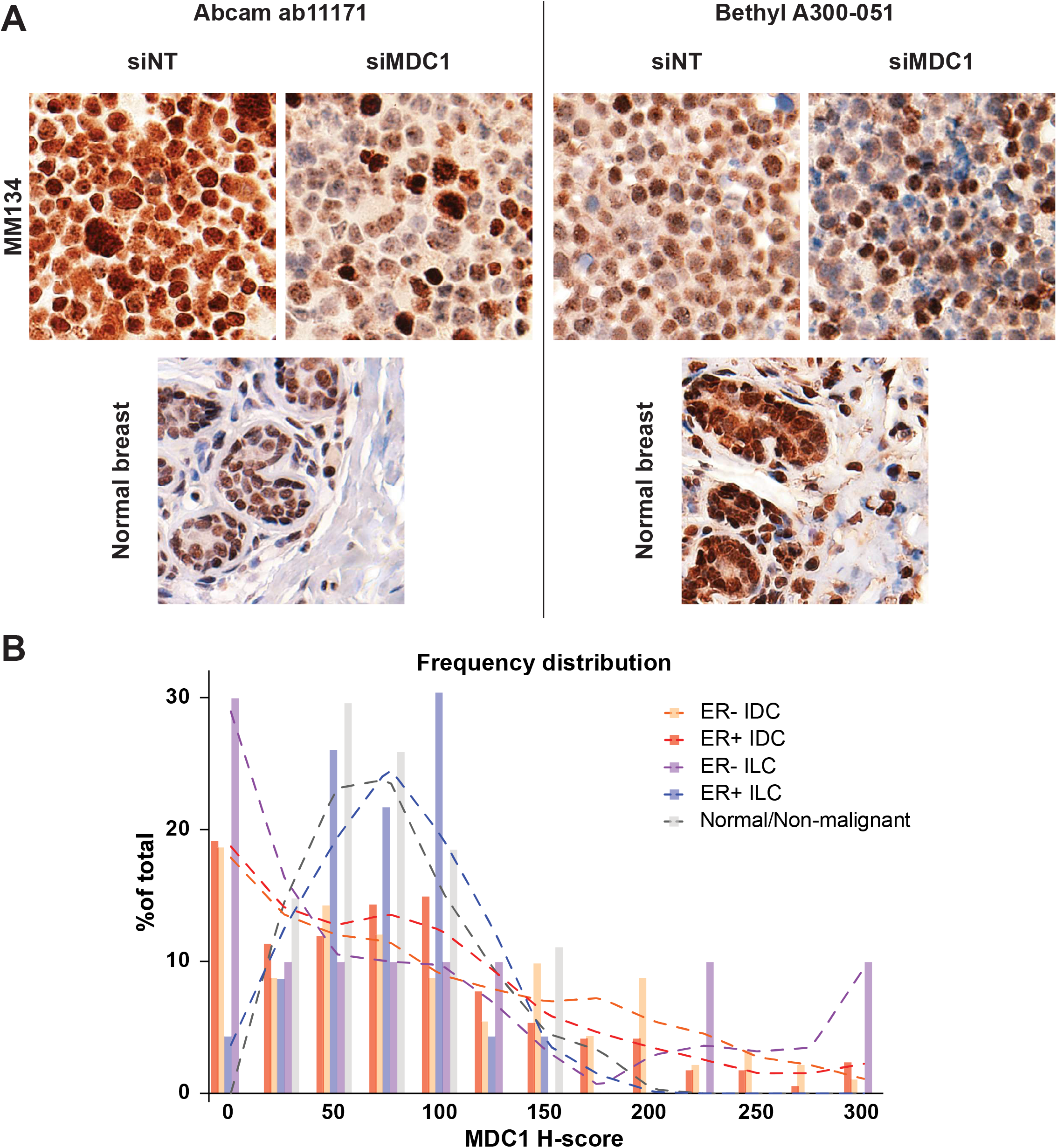
MDC1 IHC analyses in control samples and breast tumors. **(A)**, Antibodies validated by immunoblot with indications for use in IHC were tested using FFPE cell pellets from MM134 48hrs post-siRNA transfection, and using sections from normal breast tissues. Abcam 11171 was selected due to greater specificity and reduced stromal staining in tissue sections. **(B)**, Histogram of MDC1 H-scores. Dashed lines are LOWESS spline fits, and support that MDC1 H-score distribution in ER+ ILC mirrors normal tissue samples, but not other breast tumors.

## REFERENCES

1. Dossus L, Benusiglio PR. Lobular breast cancer: incidence and genetic and non-genetic risk factors. Breast Cancer Res.BioMed Central; 2015;17(1):37. PMID: 25848941

2. Ciriello G, Gatza ML, Beck AH, Wilkerson MD, Rhie SK, Pastore A, et al. Comprehensive Molecular Portraits of Invasive Lobular Breast Cancer. Cell. 2015 Oct 8;163(2):506–19. PMID: 26451490

3. Sikora MJ, Jankowitz RC, Dabbs DJ, Oesterreich S. Invasive lobular carcinoma of the breast: patient response to systemic endocrine therapy and hormone response in model systems. Steroids. Elsevier Inc.; 2013 Jun;78(6):568–575. PMID: 23178159

4. Thomas M, Kelly ED, Abraham J, Kruse M. Invasive lobular breast cancer: A review of pathogenesis, diagnosis, management, and future directions of early stage disease. Semin Oncol. 2019;46(2):121–132. PMID: 31239068

5. Colleoni M, Rotmensz N, Maisonneuve P, Mastropasqua MG, Luini A, Veronesi P, et al. Outcome of special types of luminal breast cancer. Ann Oncol Off J Eur Soc Med Oncol. 2012 Jun 29;23(6):1428–36. PMID: 22039080

6. Pestalozzi BC, Zahrieh D, Mallon E, Gusterson BA, Price KN, Gelber RD, et al. Distinct clinical and prognostic features of infiltrating lobular carcinoma of the breast: combined results of 15 International Breast Cancer Study Group clinical trials. J Clin Oncol. 2008 Jun 20;26(18):3006–14. PMID: 18458044

7. Rakha EA, El-Sayed ME, Powe DG, Green AR, Habashy H, Grainge MJ, et al. Invasive lobular carcinoma of the breast: response to hormonal therapy and outcomes. Eur J Cancer. 2008 Jan;44(1):73–83. PMID: 18035533

8. Chen Z, Yang J, Li S, Lv M, Shen Y, Wang B, et al. Invasive lobular carcinoma of the breast: A special histological type compared with invasive ductal carcinoma. PLoS One. 2017;12(9):e0182397. PMID: 28863134

9. Pan H, Gray R, Braybrooke J, Davies C, Taylor C, McGale P, et al. 20-Year Risks of Breast-Cancer Recurrence after Stopping Endocrine Therapy at 5 Years. N Engl J Med. 2017;377(19):1836–1846. PMID: 29117498

10. Metzger Filho O, Giobbie-Hurder A, Mallon E, Gusterson B, Viale G, Winer EP, et al. Relative Effectiveness of Letrozole Compared With Tamoxifen for Patients With Lobular Carcinoma in the BIG 198 Trial. J Clin Oncol. 2015 Sep 1;33(25):2772–9. PMID: 26215945

11. Knauer M, Gruber C, Dietze O, Greil R, Stöger H, Rudas M, et al. Abstract S2-06: Survival advantage of anastrozol compared to tamoxifen for lobular breast cancer in the ABCSG-8 study. Cancer Res. 2015 Apr 30;75(9 Supplement):S2-06-S2-06.

12. Sikora MJ, Cooper KL, Bahreini A, Luthra S, Wang G, Chandran UR, et al. Invasive lobular carcinoma cell lines are characterized by unique estrogen-mediated gene expression patterns and altered tamoxifen response. Cancer Res. 2014 Mar 1;74(5):1463–74. PMID: 24425047

13. Tasdemir N, Bossart EA, Li Z, Zhu L, Sikora MJ, Levine KM, et al. Comprehensive Phenotypic Characterization of Human Invasive Lobular Carcinoma Cell Lines in 2D and 3D Cultures. Cancer Res. 2018 Nov 1;78(21):6209–6222. PMID: 30228172

14. Bouaboula M, Shomali M, Cheng J, Malkova N, Sun F, Koundinya M, et al. SAR439859, an orally bioavailable selective estrogen receptor degrader (SERD) that demonstrates robust antitumor efficacy and limited cross-resistance in ER+ breast cancer [abstract]. Proc 109th Annu Meet Am Assoc Cancer Res. 2018;Chicago,IL(Philadelphia (PA): AACR):Abstract 943.

15. Guan J, Zhou W, Hafner M, Blake RA, Chalouni C, Chen IP, et al. Therapeutic Ligands Antagonize Estrogen Receptor Function by Impairing Its Mobility. Cell. 2019;178(4):949–963.e18. PMID: 31353221

16. Sreekumar S, Levine KM, Sikora MJ, Chen J, Tasdemir N, Carter D, et al. Differential Regulation and Targeting of Estrogen Receptor α Turnover in Invasive Lobular Breast Carcinoma. Endocrinology. 2020 Sep 1;161(9). PMID: 32609836

17. Sikora MJ, Jacobsen BM, Levine K, Chen J, Davidson NE, Lee A V, et al. WNT4 mediates estrogen receptor signaling and endocrine resistance in invasive lobular carcinoma cell lines. Breast Cancer Res. Breast Cancer Research; 2016 Sep 20;18(1):92. PMID: 27650553

18. Shackleford MT, Rao DM, Bordeaux EK, Hicks HM, Towers CG, Sottnik JL, et al. Estrogen Regulation of mTOR Signaling and Mitochondrial Function in Invasive Lobular Carcinoma Cell Lines Requires WNT4. Cancers (Basel). 2020 Oct 12;12(10). PMID: 33053661

19. Rao DM, Shackleford MT, Bordeaux EK, Sottnik JL, Ferguson RL, Yamamoto TM, et al. Wnt family member 4 (WNT4) and WNT3A activate cell-autonomous Wnt signaling independent of porcupine O- acyltransferase or Wnt secretion. J Biol Chem. 2019 Dec 27;294(52):19950–19966. PMID: 31740580

20. McKenna NJ, O’Malley BW. Combinatorial control of gene expression by nuclear receptors and coregulators. Cell. 2002 Feb 22;108(4):465–74. PMID: 11909518

21. Feng Q, O’Malley BW. Nuclear receptor modulation--role of coregulators in selective estrogen receptor modulator (SERM) actions. Steroids. 2014 Nov;90:39–43. PMID: 24945111

22. Han SJ, O’Malley BW. The dynamics of nuclear receptors and nuclear receptor coregulators in the pathogenesis of endometriosis. Hum Reprod Update. 2014 Jul 1;20(4):467–484. PMID: 24634322

23. Hurtado A, Holmes K a, Ross-Innes CS, Schmidt D, Carroll JS. FOXA1 is a key determinant of estrogen receptor function and endocrine response. Nat Genet. Nature Publishing Group; 2011 Jan;43(1):27–33. PMID: 21151129

24. Theodorou V, Stark R, Menon S, Carroll JS. GATA3 acts upstream of FOXA1 in mediating ESR1 binding by shaping enhancer accessibility. Genome Res. 2013 Jan 1;23(1):12–22. PMID: 23172872

25. Mohammed H, Taylor C, Brown GD, Papachristou EK, Carroll JS, D’Santos CS. Rapid immunoprecipitation mass spectrometry of endogenous proteins (RIME) for analysis of chromatin complexes. Nat Protoc. 2016 Feb;11(2):316–26. PMID: 26797456

26. Mohammed H, D’Santos C, Serandour A a., Ali HR, Brown GD, Atkins A, et al. Endogenous purification reveals GREB1 as a key estrogen receptor regulatory factor. Cell Rep. The Authors; 2013 Feb 21;3(2):342–9. PMID: 23403292

27. Jungmichel S, Stucki M. MDC1: The art of keeping things in focus. Chromosoma. 2010 Aug;119(4):337–49. PMID: 20224865

28. Coster G, Goldberg M. The cellular response to DNA damage: a focus on MDC1 and its interacting proteins. Nucleus. 2010;1(2):166–78. PMID: 21326949

29. Szklarczyk D, Gable AL, Lyon D, Junge A, Wyder S, Huerta-Cepas J, et al. STRING v11: protein-protein association networks with increased coverage, supporting functional discovery in genome-wide experimental datasets. Nucleic Acids Res. 2019;47(D1):D607–D613. PMID: 30476243

30. Martin L-A, Ribas R, Simigdala N, Schuster E, Pancholi S, Tenev T, et al. Discovery of naturally occurring ESR1 mutations in breast cancer cell lines modelling endocrine resistance. Nat Commun. 2017 Nov 30;8(1):1865. PMID: 29192207

31. Stewart GS, Wang B, Bignell CR, Taylor AMR, Elledge SJ. MDC1 is a mediator of the mammalian DNA damage checkpoint. Nature. 2003 Feb 27;421(6926):961–6. PMID: 12607005

32. Goldberg M, Stucki M, Falck J, D’Amours D, Rahman D, Pappin D, et al. MDC1 is required for the intra-S-phase DNA damage checkpoint. Nature. 2003 Feb 27;421(6926):952–6. PMID: 12607003

33. Liberzon A, Birger C, Thorvaldsdóttir H, Ghandi M, Mesirov JP, Tamayo P. The Molecular Signatures Database Hallmark Gene Set Collection. Cell Syst. 2015;1(6):417–425. PMID: 26771021

34. Schiewer MJ, Knudsen KE. Linking DNA Damage and Hormone Signaling Pathways in Cancer. Trends Endocrinol Metab. Elsevier Ltd; 2016 Apr;27(4):216–225. PMID: 26944914

35. Imrichová H, Hulselmans G, Atak ZK, Potier D, Aerts S. i-cisTarget 2015 update: generalized cis-regulatory enrichment analysis in human, mouse and fly. Nucleic Acids Res. 2015 Jul 1;43(W1):W57–64. PMID: 25925574

36. Ruff SE, Logan SK, Garabedian MJ, Huang TT. Roles for MDC1 in cancer development and treatment. DNA Repair (Amst). 2020 Nov;95(July):102948. PMID: 32866776

37. Patel AN, Goyal S, Wu H, Schiff D, Moran MS, Haffty BG. Mediator of DNA damage checkpoint protein 1 (MDC1) expression as a prognostic marker for nodal recurrence in early-stage breast cancer patients treated with breast-conserving surgery and radiation therapy. Breast Cancer Res Treat. 2011 Apr;126(3):601–7. PMID: 20521098

38. Bartkova J, Horejsí Z, Sehested M, Nesland JM, Rajpert-De Meyts E, Skakkebaek NE, et al. DNA damage response mediators MDC1 and 53BP1: constitutive activation and aberrant loss in breast and lung cancer, but not in testicular germ cell tumours. Oncogene. 2007 Nov 22;26(53):7414–22. PMID: 17546051

39. Zou R, Zhong X, Wang C, Sun H, Wang S, Lin L, et al. MDC1 Enhances Estrogen Receptor-mediated Transactivation and Contributes to Breast Cancer Suppression. Int J Biol Sci. 2015;11(9):992–1005. PMID: 26221067

40. Cerami E, Gao J, Dogrusoz U, Gross BE, Sumer SO, Aksoy BA, et al. The cBio cancer genomics portal: an open platform for exploring multidimensional cancer genomics data. Cancer Discov. 2012 May;2(5):401–4. PMID: 22588877

41. Solier S, Pommier Y. MDC1 cleavage by caspase-3: a novel mechanism for inactivating the DNA damage response during apoptosis. Cancer Res. 2011 Feb 1;71(3):906–13. PMID: 21148072

42. Stires H, Heckler MM, Fu X, Li Z, Grasso CS, Quist MJ, et al. Integrated molecular analysis of Tamoxifen-resistant invasive lobular breast cancer cells identifies MAPK and GRM/mGluR signaling as therapeutic vulnerabilities. Mol Cell Endocrinol. 2018;471:105–117. PMID: 28935545

43. Riggins RB, Lan JP-J, Zhu Y, Klimach U, Zwart A, Cavalli LR, et al. ERRgamma mediates tamoxifen resistance in novel models of invasive lobular breast cancer. Cancer Res. 2008/11/01. 2008 Nov 1;68(21):8908–17. PMID: 18974135

44. Ozaki T, Nagase T, Ichimiya S, Seki N, Ohiri M, Nomura N, et al. NFBD1/KIAA0170 is a novel nuclear transcriptional transactivator with BRCT domain. DNA Cell Biol. 2000 Aug;19(8):475–85. PMID: 10975465

45. Wan C, Borgeson B, Phanse S, Tu F, Drew K, Clark G, et al. Panorama of ancient metazoan macromolecular complexes. Nature. 2015 Sep 17;525(7569):339–44. PMID: 26344197

46. Jung SY, Malovannaya A, Wei J, O’Malley BW, Qin J. Proteomic analysis of steady-state nuclear hormone receptor coactivator complexes. Mol Endocrinol. 2005 Oct;19(10):2451–65. PMID: 16051665

47. Woods NT, Mesquita RD, Sweet M, Carvalho MA, Li X, Liu Y, et al. Charting the landscape of tandem BRCT domain-mediated protein interactions. Sci Signal. 2012 Sep 18;5(242):rs6. PMID: 22990118

48. Johnson AB, O’Malley BW. Steroid receptor coactivators 1, 2, and 3: critical regulators of nuclear receptor activity and steroid receptor modulator (SRM)-based cancer therapy. Mol Cell Endocrinol. 2012 Jan 30;348(2):430–9. PMID: 21664237

49. Periyasamy M, Patel H, Lai C-F, Nguyen VTM, Nevedomskaya E, Harrod A, et al. APOBEC3B-Mediated Cytidine Deamination Is Required for Estrogen Receptor Action in Breast Cancer. Cell Rep. 2015 Oct 6;13(1):108–121. PMID: 26411678

50. Wang C, Sun H, Zou R, Zhou T, Wang S, Sun S, et al. MDC1 functionally identified as an androgen receptor co-activator participates in suppression of prostate cancer. Nucleic Acids Res. 2015 May 26;43(10):4893–908. PMID: 25934801

51. Li Z, Shao C, Kong Y, Carlock C, Ahmad N, Liu X. DNA Damage Response-Independent Role for MDC1 in Maintaining Genomic Stability. Mol Cell Biol. 2017 May 1;37(9):1–17. PMID: 28193847

52. Lou Z, Minter-Dykhouse K, Franco S, Gostissa M, Rivera MA, Celeste A, et al. MDC1 maintains genomic stability by participating in the amplification of ATM-dependent DNA damage signals. Mol Cell. 2006 Jan 20;21(2):187–200. PMID: 16427009

53. Lee J-H, Park S-J, Jeong S-Y, Kim M-J, Jun S, Lee H-S, et al. MicroRNA-22 Suppresses DNA Repair and Promotes Genomic Instability through Targeting of MDC1. Cancer Res. 2015 Apr 1;75(7):1298–310. PMID: 25627978

54. Minter-Dykhouse K, Ward I, Huen MSY, Chen J, Lou Z. Distinct versus overlapping functions of MDC1 and 53BP1 in DNA damage response and tumorigenesis. J Cell Biol. 2008;181(5):727–735. PMID: 18504301

55. Baker LA, Baker LA, Holliday H, Holliday H, Roden D, Roden D, et al. Proteogenomic analysis of Inhibitor of Differentiation 4 (ID4) in basal-like breast cancer. Breast Cancer Res. Breast Cancer Research; 2020;22(1):1–18. PMID: 32527287

56. Luo K, Zhang H, Wang L, Yuan J, Lou Z. Sumoylation of MDC1 is important for proper DNA damage response. EMBO J. 2012 Jun 29;31(13):3008–19. PMID: 22635276

57. Watanabe S, Watanabe K, Akimov V, Bartkova J, Blagoev B, Lukas J, et al. JMJD1C demethylates MDC1 to regulate the RNF8 and BRCA1-mediated chromatin response to DNA breaks. Nat Struct Mol Biol. Nature Publishing Group; 2013 Dec;20(12):1425–33. PMID: 24240613

58. Salguero I, Belotserkovskaya R, Coates J, Sczaniecka-Clift M, Demir M, Jhujh S, et al. MDC1 PST-repeat region promotes histone H2AX-independent chromatin association and DNA damage tolerance. Nat Commun. Springer US; 2019 Nov 15;10(1):5191. PMID: 31729360

59. Akbani R, Ng PKS, Werner HMJ, Shahmoradgoli M, Zhang F, Ju Z, et al. A pan-cancer proteomic perspective on The Cancer Genome Atlas. Nat Commun. 2014 May 29;5:3887. PMID: 24871328

60. Riba LA, Russell T, Alapati A, Davis RB, James TA. Characterizing Response to Neoadjuvant Chemotherapy in Invasive Lobular Breast Carcinoma. J Surg Res. Elsevier Inc; 2019 Jan;233:436–443. PMID: 30502283

61. Tadros AB, Wen HY, Morrow M. Breast Cancers of Special Histologic Subtypes Are Biologically Diverse. Ann Surg Oncol. 2018 Oct;25(11):3158–3164. PMID: 30094484

62. Loibl S, Volz C, Mau C, Blohmer J-U, Costa SD, Eidtmann H, et al. Response and prognosis after neoadjuvant chemotherapy in 1,051 patients with infiltrating lobular breast carcinoma. Breast Cancer Res Treat. 2014 Feb;144(1):153–62. PMID: 24504379

63. Truin W, Voogd AC, Vreugdenhil G, van der Heiden-van der Loo M, Siesling S, Roumen RM. Effect of adjuvant chemotherapy in postmenopausal patients with invasive ductal versus lobular breast cancer. Ann Oncol. 2012;23(11):2859–2865. PMID: 22745216

64. Jambal P, Badtke MM, Harrell JC, Borges VF, Post MD, Sollender GE, et al. Estrogen switches pure mucinous breast cancer to invasive lobular carcinoma with mucinous features. Breast Cancer Res Treat. 2013 Jan;137(2):431–48. PMID: 23247610

65. Sikora MJ, Johnson MD, Lee A V, Oesterreich S. Endocrine Response Phenotypes Are Altered by Charcoal-Stripped Serum Variability. Endocrinology. 2016 Oct 26;157(10):3760–3766. PMID: 27459541

66. Mellacheruvu D, Wright Z, Couzens AL, Lambert J-P, St-Denis NA, Li T, et al. The CRAPome: a contaminant repository for affinity purification-mass spectrometry data. Nat Methods. 2013 Aug;10(8):730–6. PMID: 23921808

67. Ivashkevich AN, Martin OA, Smith AJ, Redon CE, Bonner WM, Martin RF, et al. γH2AX foci as a measure of DNA damage: a computational approach to automatic analysis. Mutat Res. 2011 Jun 3;711(1- 2):49–60. PMID: 21216255

68. Bushnell B. BBMap [Internet].

69. Dobin A, Davis CA, Schlesinger F, Drenkow J, Zaleski C, Jha S, et al. STAR: ultrafast universal RNA-seq aligner. Bioinformatics. 2013 Jan 1;29(1):15–21. PMID: 23104886

70. Robinson MD, McCarthy DJ, Smyth GK. edgeR: a Bioconductor package for differential expression analysis of digital gene expression data. Bioinformatics. 2010 Jan 1;26(1): 139–40. PMID: 19910308

71. Ritchie ME, Phipson B, Wu D, Hu Y, Law CW, Shi W, et al. limma powers differential expression analyses for RNA-sequencing and microarray studies. Nucleic Acids Res. 2015 Apr 20;43(7):e47. PMID: 25605792

72. Yu G, Wang L-G, Han Y, He Q-Y. clusterProfiler: an R package for comparing biological themes among gene clusters. OMICS. 2012 May;16(5):284–7. PMID: 22455463

73. Aibar S, González-Blas CB, Moerman T, Huynh-Thu VA, Imrichova H, Hulselmans G, et al. SCENIC: single-cell regulatory network inference and clustering. Nat Methods. 2017 Nov;14(11):1083–1086. PMID: 28991892

74. Howe EA, Sinha R, Schlauch D, Quackenbush J. RNA-Seq analysis in MeV. Bioinformatics. 2011 Nov 15;27(22):3209–10. PMID: 21976420

