## Supplemental Files 1-11 for "Mediator of DNA damage checkpoint 1 (MDC1) is a novel estrogen receptor co-regulator in invasive lobular carcinoma of the breast": Supplemental File 9 - Active Motif RIME protocol.pdf

### RIME CELL FIXATION PROTOCOL

1. Fix 5 x $10^7$ to $10^8$ cells for each cell population to be tested. To fix, add 1/10 volume of freshly-prepared Formaldehyde Solution* (see Reagents below) to the existing media. Do NOT remove existing media. For example, to a flask containing 10 ml of media, add 1 ml of Formaldehyde Solution. Cap and agitate for **8** minutes at room temperature.
2. Stop the fixation by adding 1/20 volume Glycine Solution* to the existing media in each container. For example, if the flask from Step 1 now contains 11 ml, add 0.55 ml 2.5 M glycine. Let sit at room temperature for 5 minutes. After the glycine incubation, if the cells are adherent, scrape them thoroughly from the culture surface.
3. Wash cells by transferring contents of each container to a conical tube (15 ml or 50 ml tube, depending on the volume). Keep samples on ice for the remainder of the procedure. Centrifuge tubes at 800 x *g* in a refrigerated centrifuge for 10 minutes to pellet the cells. Remove the supernatant and re-suspend cells in 10 ml chilled PBS-Igepal* per tube by pipetting up and down. If cells from any one population are contained in multiple centrifuge tubes, combine them at this step.
4. Centrifuge tubes again as before to pellet the cells. Remove the supernatant, then add 10 ml chilled PBS-Igepal* to each tube. Add 100 µl PMSF (100 mM in ethanol*; final concentration will be 1 mM) to each tube and pipet up and down to re-suspend the cells.
5. Centrifuge tubes a third time to pellet the cells, and carefully remove supernatant completely from cell pellets.
6. Snap-freeze cell pellets on dry ice and store at -80°C.
7. Ship cells on dry ice to Active Motif. Follow the instructions shown on our Sample Submission Form, which can be downloaded at: www.activemotif.com/sample-submission.

#### Reagents*

| | Final concentration | Per 10 ml |
|---|---|---|
| 1. Formaldehyde Solution (to be prepared fresh before use): | | |
| 16% Methanol Free Formaldehyde (*e.g.* ThermoFisher #28908) | 11% | 6.9 ml |
| 5 M NaCl | 0.1 M | 200 µl |
| 0.5 M EDTA, pH 8.0 | 1 mM | 20 µl |
| 1 M HEPES, pH 7.9 | 50 mM | 500 µl |
| $H_2O$ | | 2.4 ml |
| (Note: NaCl, EDTA, and HEPES should be molecular biology grade.) | | |

| 2. Glycine Solution | | Per 10 ml |
|---|---|---|
| Glycine, MW 75 (*e.g.* Sigma #G-7403) | 2.5 M | 1.88 g |
| $H_2O$ | | to 10 ml |

| 3. PBS-Igepal | | Per 100 ml |
|---|---|---|
| PBS, pH 7.4 (*e.g.* ThermoFisher #10010023) | ~1X | 100 ml |
| 100% Igepal CA-630 (*e.g.* Sigma #I-8896) | 0.5% | 0.5 ml |

4. PMSF (e.g. Sigma #P-7626)

Prepare at 100 mM in ethanol and store at -20°C.
(Note: PMSF Phenylmethanesulfonyl fluoride.)
