## Supplemental Files 1-11 for "Mediator of DNA damage checkpoint 1 (MDC1) is a novel estrogen receptor co-regulator in invasive lobular carcinoma of the breast": Supplemental File 10 - RIME QC Reports for ERa.pdf

Enabling Epigenetics Research

### ERa RIME Report

Project: 3317UCol
Investigator: Matthew Sikora

**Experiment**: RIME (**R**apid **I**mmunoprecipitation **M**ass Spectrometry of **E**ndogenous Proteins) was carried out in duplicate using an antibody against ERa (Santa Cruz, Cat. sc-543) and 150 µg of chromatin from MM134 cells to identify proteins that interact with ERa using mass spectrometry.

**Summary**: Mass spectrometry of the immunoprecipitated protein complexes resulted in good (34-36%) coverage of the bait protein ERa, and has met our minimum requirement of 10% coverage needed for a passing experiment according to the guidelines below.

- >40% coverage =great
- 20%-39% coverage =good
- 10%-19% coverage = fair
- <10% coverage = failed

You will find an associated raw data file representing results from the experiment from a human protein database. This will portray the proteins detected by mass spectrometry and their respective amounts. Upon filtering the experimental reaction data against the negative control IgG reaction data there were a number of unique proteins pulled down along with ERa.

#### Detection of Target (bait) Protein:

| Sequence Coverage | Protein | Prob | %Cov | Bio Sample |
|---|---|---|---|---|
| | Estrogen receptor OS=Homo sapiens ... | 100% | 34% | MM134+Veh-R1 |
| | Estrogen receptor OS=Homo sapiens ... | 100% | 34% | MM134+Veh-R2 |
| | Estrogen receptor OS=Homo sapiens ... | 100% | 34% | MM134+4OHT-R1 |
| | Estrogen receptor OS=Homo sapiens ... | 100% | 36% | MM134+4OHT-R2 |

Regions shown in yellow represent peptides detected by mass spectrometry for the bait protein (ERa). Green bars represent those for which post-translational modifications were detected.

### ERa RIME Report

Project: 3317UCol
Investigator: Matthew Sikora

**Experiment**: RIME (**R**apid **I**mmunoprecipitation **M**ass Spectrometry of **E**ndogenous Proteins) was carried out in duplicate using an antibody against ERa (Santa Cruz, Cat. sc-543) and 150 µg of chromatin from SUM44PE cells to identify proteins that interact with ERa using mass spectrometry.

**Summary**: Mass spectrometry of the immunoprecipitated protein complexes resulted in good (16-25%) coverage of the bait protein ERa, and has met our minimum requirement of 10% coverage needed for a passing experiment according to the guidelines below.

- >40% coverage =great
- 20%-39% coverage =good
- 10%-19% coverage = fair
- <10% coverage = failed.

You will find an associated raw data file representing results from the experiment from a human protein database. This will portray the proteins detected by mass spectrometry and their respective amounts. Upon filtering the experimental reaction data against the negative control IgG reaction data there were a number of unique proteins pulled down along with ERa.

**Detection of Target (bait) Protein:**

| Sequence Coverage | Protein | Prob | %Cov | Bio Sample |
|---|---|---|---|---|
| | Estrogen receptor OS=Homo sapiens ... | 100% | 16% | 44PE+Veh-R1 |
| | Estrogen receptor OS=Homo sapiens ... | 100% | 24% | 44PE+Veh-R2 |
| | Estrogen receptor OS=Homo sapiens ... | 100% | 25% | 44PE+4OHT-R1 |
| | Estrogen receptor OS=Homo sapiens ... | 100% | 21% | 44PE+4OHT-R2 |

Regions shown in yellow represent peptides detected by mass spectrometry for the bait protein (ERa). Green bars represent those for which post-translational modifications were detected.

### ERa RIME Report

Project: 3317UCol
Investigator: Matthew Sikora

**Experiment**: RIME (**R**apid **I**mmunoprecipitation **M**ass Spectrometry of **E**ndogenous Proteins) was carried out in duplicate using an antibody against ERa (Santa Cruz, Cat. sc-543) and 100 µg of chromatin from BCK4 cells to identify proteins that interact with ERa using mass spectrometry.

**Summary**: Mass spectrometry of the immunoprecipitated protein complexes resulted in fair (18-21%) coverage of the bait protein ERa, and has met our minimum requirement of 10% coverage needed for a passing experiment according to the guidelines below.

- >40% coverage =great
- 20%-39% coverage =good
- 10%-19% coverage = fair
- <10% coverage = failed

You will find an associated raw data file representing results from the experiment from a human protein database. This will portray the proteins detected by mass spectrometry and their respective amounts. Upon filtering the experimental reaction data against the negative control IgG reaction data there were a number of unique proteins pulled down along with ERa.

#### Detection of Target (bait) Protein:

| Sequence Coverage | Protein | Prob | %Cov | Bio Sample |
|---|---|---|---|---|
| | Estrogen receptor OS=Homo sapiens ... | 100% | 20% | BCK4+Veh-R1 |
| | Estrogen receptor OS=Homo sapiens ... | 100% | 18% | BCK4+Veh-R2 |
| | Estrogen receptor OS=Homo sapiens ... | 100% | 21% | BCK4+4OHT-R1 |
| | Estrogen receptor OS=Homo sapiens ... | 100% | 19% | BCK4+4OHT-R2 |

Regions shown in yellow represent peptides detected by mass spectrometry for the bait protein (ERa). Green bars represent those for which post-translational modifications were detected.
